# Linking brain-heart interactions to emotional arousal in immersive virtual reality

**DOI:** 10.1101/2024.01.04.574165

**Authors:** A. Fourcade, F. Klotzsche, S. M. Hofmann, A. Mariola, V. V. Nikulin, A. Villringer, M. Gaebler

## Abstract

The subjective experience of emotions is linked to the contextualized perception and appraisal of changes in bodily (e.g., heart) activity. Increased emotional arousal has been related to attenuated high-frequency heart rate variability (HF-HRV), lower EEG parieto-occipital alpha power, and higher heartbeat-evoked potential (HEP) amplitudes. We studied emotional arousal-related brain-heart interactions using immersive virtual reality (VR) for naturalistic yet controlled emotion induction. 29 healthy adults (13 women, age: 26±3) completed a VR experience that included rollercoasters while EEG and ECG were recorded. Continuous emotional arousal ratings were collected during a video replay immediately after. We analyzed emotional arousal-related changes in HF-HRV as well as in BHIs using HEPs. Additionally, we used the oscillatory information in the ECG and the EEG to model the directional information flows between the brain and heart activity.

We found that higher emotional arousal was associated with lower HEP amplitudes in a left fronto-central electrode cluster. While parasympathetic modulation of the heart (HF-HRV) and parieto-occipital EEG alpha power were reduced during higher emotional arousal, there was no evidence for the hypothesized emotional arousal-related changes in bidirectional information flow between them. Whole-brain exploratory analyses in additional EEG (delta, theta, alpha, beta and gamma) and HRV (low-frequency, LF, and HF) frequency bands revealed a temporo-occipital cluster, in which higher emotional arousal was linked to decreased brain-to-heart (i.e., gamma→HF-HRV) and increased heart-to-brain (i.e., LF-HRV→gamma) information flow. Our results confirm previous findings from less naturalistic experiments and suggest a link between emotional arousal and brain-heart interactions in temporo-occipital gamma power.

## Introduction

Emotions result from internal or external events that are appraised as relevant to an organism’s well-being (Gross, 1998; Rottenberg & Gross, 2003). Emotions have subjective and physiological components (Mauss & Robinson, 2009), among others (e.g., expressive; Sander et al., 2005). We aimed to link subjective emotional experience to physiological activity not only in the brain but also in the heart and their interaction under naturalistic stimulation using immersive virtual reality.

The subjective component of emotions or - more generally - of affect is a basic property of human experience (Wundt, 1897) or consciousness (Barrett, 2016), and it is linked to the contextualized perception and appraisal of bodily changes (James, 1884; Lange, 1885; Cannon, 1927; Bard, 1928; Schachter & Singer, 1962). While theories like James-Lange (James, 1884; Lange, 1885), Cannon-Bard (Cannon, 1927; Bard, 1928), or Schachter-Singer (Schachter & Singer, 1962) emphasize different temporal order and mutual importance of peripheral physiological reactions and (brain-centric) cognitive-evaluative components, all include a link between bodily processes and the subjective experience of emotions. In more recent terms, interoceptive information (e.g., autonomic, visceral) about the physiological state of the internal body (Craig, 2002; Barrett, 2016) is combined with exteroceptive information (e.g., auditory, visual) about the physical state of the external world. The subjective experience of affect is often described using the two continuous dimensions or “core affects” of valence and *emotional* arousal (Russell, 2003): Valence refers to how pleasant or unpleasant, and emotional arousal refers to how intense the experience is (Barrett & Russell, 1999).

Sweaty palms, increased heart rate, and the throat tightening at the sight of a snake are examples of physiological responses connected to emotions. These peripheral physiological responses are regulated by the autonomic nervous system (ANS), with activity of the parasympathetic branch being linked to a “rest-and-digest’’ mode and activity of the sympathetic branch to the “flight-or-fight” response (McCorry, 2007). Focusing on the cardiovascular system, the two branches can modulate the activity of the sinoatrial node in the heart and change the heart rate (Levy, 1971). These influences have different temporal dynamics, with sympathetic regulation being relatively slow (at the order of seconds) and parasympathetic regulation being relatively fast (at the order of milliseconds; Warner & Cox, 1962; Levy & Martin, 1984). Isolating the sympathetic influence on the heart has been difficult (Goldstein et al., 2011; Reyes del Paso et al., 2013; Shaffer et al., 2014). However, autonomic blocking agents (Akselrod et al., 1981; Ali-Melkkilä et al., 1991), cardiac denervation (e.g., vagotomy; Hedman et al., 1995) and body posture manipulations (Pomeranz et al., 1985) have shown that the parasympathetic influence, or vagal cardioregulation, can be quantified by high-frequency heart rate variability (HF-HRV; Task Force, 1996), that is fluctuations of the time between heartbeats (i.e., the interbeat interval, IBI) in 0.15 and 0.4 Hz frequency range and that low frequency HRV (LF-HRV; 0.04 and 0.15 Hz) reflects both sympathetic and parasympathetic influences on the heart. ANS activity has been linked to emotion, but specific patterns that are exclusively associated with particular emotion categories could not be identified (Kreibig, 2010; Siegel et al., 2018). For the dimensional core affects of emotional arousal and valence, changes in ANS activity appear more consistently in the literature; unpleasant (i.e., negatively valenced) stimuli (e.g., movie, music) were associated with a decrease in heart rate (Palomba et al., 2000; Sammler et al., 2007) and higher emotional arousal was linked to decreased HF-HRV (Valenza et al., 2012; Luft & Bhattacharya, 2015; Hildebrandt et al., 2016), suggesting a decreased parasympathetic influence on the heart during higher emotional arousal.

The integration of external (exteroceptive) and internal (interoceptive) signals (e.g., from the heart), which gives rise to emotional experiences, takes place in the brain (Craig, 2009; Seth, 2013). A major route of information flow between the body and the brain is the vagus nerve, which transmits sensory signals from the heart (through the dorsal root and stellate ganglia; Dusi & Ardell, 2020) to the brainstem, where they result in reflexes (e.g., the baroreflex) but are also relayed to the cerebrum and cerebellum. Reversely, preganglionic autonomic motor neurons are also innervated by the vagus nerve and controlled by the brain; in particular by the central autonomic network (CAN), which includes the hypothalamus, amygdala, anterior cingulate, insula and medial prefrontal cortex (Benarroch, 1993; Thayer & Lane, 2000, 2009). These brain regions are also consistently reported in neuroimaging studies of emotions (Dalgleish, 2004; Lindquist et al., 2012). Recent evidence from rodents suggests that particularly insula, amygdala, and prefrontal cortex but also brainstem circuits integrate cardiac and vagal activity into information processing, influencing emotion perception and adaptive behavior (Gehrlach et al., 2019; Hsueh et al., 2023; Klein et al., 2021; Signoret-Genest et al., 2023; Okonogi et al., 2024). Optogenetic stimulation of the heart (Hsueh et al., 2023) and individual brain regions (Signoret-Genest et al., 2023; Klein et al., 2021), as well as vagus-nerve surgery and stimulation (Okonogi et al., 2024), thereby provide causal evidence for the bidirectional information flow between heart and the brain and its role for affective processing.

The activation in the heart and the brain can be electrophysiologically recorded using ECG and EEG, respectively. Both signals contain oscillatory and aperiodic components (heart: Babloyantz & Destexhe, 1988; brain: Buzsáki et al., 2013; He, 2014). In the ECG signal, oscillations in HRV are typically decomposed into LF- and HF-HRV, while in the EEG signal, five frequency bands are commonly defined: delta (δ; 0.3–4 Hz), theta (θ; 4–8 Hz), alpha (α; 8–13 Hz), beta (β; 13–30 Hz) and gamma (γ; 30–45 Hz). Both ECG and EEG oscillations are modulated by non-invasive vagus nerve stimulation (Machetanz et al., 2021), which supports the results from invasive manipulations in animals (see above).

Various functions, from more fundamental sensory and motor to higher cognitive processes, have been ascribed to different types of regionally distributed neural oscillations (Buzsaki, 2006). In particular, the parieto-occipital alpha rhythm is the dominant EEG rhythm in awake adults with eyes closed (Berger, 1929), where it varies with vigilance (Olbrich et al., 2009). Its physiological bases are large-scale synchronization of neuronal activity (Buzsaki, 2006) and metabolic deactivation (Moosmann et al., 2003). Psychophysiologically, parieto-occipital alpha power is correlated to emotional arousal (Koelstra et al., 2012; Luft & Bhattacharya, 2015; Hofmann, Klotzsche, Mariola et al., 2021), as well as to attentional processes (Klimesch, 2012; van Diepen et al., 2019) which help prioritize and select sensory inputs (Treisman, 1969; Driver, 2001). Particularly bottom-up or stimulus-driven attention is assumed to direct perception towards a subset of salient external stimuli (Egeth & Yantis, 1997).

Besides sharing an electrophysiological pattern, affect and attention have been psychologically linked. On the one hand, emotional stimuli capture attentional focus (Nummenmaa et al., 2006, 2009) and their processing is prioritized over non-emotional stimuli (Okon-Singer et al., 2013). On the other hand, changes in affective states (and particularly the level of emotional arousal) alter attention-related changes in salience also for non-emotional stimuli (Sutherland & Mather, 2018).

In sum, as activity in the heart and the brain play important roles for affective experiences, their multimodal or joint analysis promises to capture the physiology of human experience more comprehensively than the analysis of either of the modalities alone (e.g., Raimondo et al., 2017). Emotion-related brain-heart interactions (BHIs), typically measured through electrophysiological recordings, have been investigated using different methodologies: for instance, by means of event-related (e.g., heartbeat-evoked potentials [HEP]; Luft & Bhattacharya, 2015) and oscillatory analyses (e.g., directional functional BHI: Candia-Rivera et al., 2022).

The HEP is an event-related potential (ERP) component that can be observed when the EEG signal is time-locked to the R-peaks in the ECG (Schandry et al., 1986). The HEP is taken to reflect the cortical processing of the heartbeat (Park & Blanke, 2019) and its amplitude is modulated by interoceptive (Pollatos et al., 2016) and emotional processing. For example, Luft and Bhattacharya (2015) manipulated emotional arousal through videos and music (NB: without assessing subjective experience) and found higher HEP amplitudes during blocks of higher (HA) compared to lower emotional arousal (LA) - particularly over right parietal electrodes (P6 and P8) from 0.38 to 0.46 s after the R-peak. Higher HEP amplitude has been interpreted as reflecting a shift of attentional focus from external stimuli to internal bodily states (García-Cordero et al., 2017; Villena-González et al., 2017; Petzschner et al., 2019; Zaccaro et al., 2022).

Directional functional BHI can be computed by jointly analyzing oscillations in EEG and ECG data, for example with the synthetic data generation (SDG) model (Catrambone et al., 2019). Based on the bidirectional communication loop in which ongoing HRV (as a proxy for ANS activity) modulates EEG (as a proxy for CNS activity) and, in turn, ongoing EEG activity modulates HRV, the SDG combines generative models for the EEG and the heartbeat. Using this approach, Candia-Rivera and colleagues (2022) recently found that the ascending information flow from the vagal cardioregulation (i.e., HF-HRV) to the EEG oscillations in the delta, theta, and gamma bands correlated with emotional arousal ratings, particularly in frontal and parieto-occipital electrodes. Furthermore, higher ascending heart-to-brain information flow preceded lower descending brain-to-heart information flow and these emotional arousal-related changes in BHI were associated with HF-HRV but not LF-HRV (Candia-Rivera et al., 2022).

Besides advances in the multimodal analysis of physiological signals, affective neuroscience is currently experiencing advances in emotion elicitation. Past research has extensively used static images repeatedly presented in trial-based designs, often creating an artificial and discontinuous experience (Bridwell et al., 2018; Huk et al., 2018). However, affective phenomena, along with their associated physiological responses, are varying on different continuous time scales; for example, emotional arousal can fluctuate on the order of minutes (Kuppens et al., 2010) or seconds (Mikutta et al., 2012). Dynamic stimuli that capture such naturalistic fluctuations in affective experience could be videos (Samide et al., 2020) or even emotional stimuli in everyday life (Grosse Rueschkamp et al., 2019). While the former remain relatively artificial (e.g., non-interactive), the latter are difficult to experimentally control (e.g., comprehensively measure). Immersive Virtual Reality (VR; wherein the user is surrounded by the virtual environment) presents a trade-off between naturalism and experimental control (Bohil et al., 2011) and offers several advantages: It enables (1) the creation and presentation of computer-generated scenarios that are contextually rich and engaging (Diemer et al., 2015), (2) naturalistic elicitation of specific psychological states (which may be particularly relevant for affective states; Baumgartner et al., 2006; Riva et al., 2007; Dores et al., 2014; Meuleman & Rudrauf, 2021), and (3) the recording of multimodal measurements (e.g., subjective ratings, movements, heart and brain signals) - with more experimental control (and less noise) than in real-life assessments (Meuleman & Rudrauf, 2021).

Recently, we used an emotionally arousing VR experience (including two rollercoasters with a break in-between) while EEG and ECG were recorded and emotional arousal was continuously rated (Hofmann, Klotzsche, Mariola et al., 2021). Using spatial filtering and non-linear machine learning for the analysis of the EEG data, we could decode subjective emotional arousal from parieto-occipital alpha power. Given the importance of cardiac activity and the heart-brain axis for affective experience and emotional arousal, we now included the ECG into the analysis. We thereby expected the joint analysis to offer insights about the physiology of emotional arousal that are not accessible when investigating EEG signals alone. In particular, we investigated the role of vagal cardioregulation for the EEG alpha power reduction during HA compared to LA, and our hypotheses were:

1. Cardiac activity and vagal cardioregulation (indexed by IBIs and HF-HRV, respectively) differ between states of HA and LA.
2. HEP amplitudes differ between states of HA and LA, particularly around 400 ms after a heartbeat and in right temporo-parietal electrodes (based on Luft & Bhattacharya, 2015).
3. Emotional arousal-related changes in parieto-occipital alpha oscillations (Hofmann, Klotzsche, Mariola et al., 2021) are related to changes in heart activity (i.e., HF-HRV).

## Materials and Methods

### 1. Participants

Participants were recruited via the participant database at the Berlin School of Mind and Brain (an adaptation of ORSEE; Greiner, 2015). Inclusion criteria were right-handedness, normal or corrected-to-normal vision, proficiency in German, no (self-reported) psychiatric, or neurological diagnoses in the past 10 years, and less than 3 hr of experience with VR. Participants were requested not to drink coffee or other stimulants 1 hr before coming to the lab. The experiment took ∼2.5 hr, and participants were reimbursed with 9 € per hour. They signed informed consent before their participation, and the study was approved by the Ethics Committee of the Department of Psychology at the Humboldt-Universität zu Berlin. Forty-five healthy young adults (22 men, mean age: 23±4, range: 20-32 years) came to the lab. Data from eleven participants needed to be discarded due to technical problems (*n* = 5), electrode malfunctioning (EEG: *n* = 1, ECG: *n* = 3), discontinuation of the experiment (*n* = 1) or violation of inclusion criteria (*n* = 1), so that data from 34 participants were processed. After EEG quality assurance (details below), data from 29 participants (16 men, mean age: 26±3, range: 20-31 years) entered the analyses.

### 2. Materials

*EEG* (sampled at 500 Hz, hardware-based low-pass filter at 131 Hz) was recorded with 30 active Ag/AgCl electrodes attached according to the international 10-20 system (actiCap and LiveAmp, Brain Products GmbH, Germany) and referenced to electrode FCz. Two additional electrodes captured eye movements (electrooculography) and were placed below and next to the right eye. *ECG* was synchronously recorded (sampled at 500 Hz) with additional electrodes in a bipolar Lead I configuration. A grounding electrode was placed on the right clavicle and two passive lead electrodes were positioned bilaterally on the participant’s torso (lower rib cage). In addition, skin conductance was recorded at the index and annulary fingers of the left hand. As the focus of this manuscript are (particularly oscillatory) brain-heart or EEG-ECG interactions, skin conductance data was not considered here.

*VR setup*: A HTC Vive HMD (HTC, Taiwan) with headphones was attached on top of the EEG cap using cushions to avoid pressure artifacts. The VR experience was commercially available on Steam (“Russian VR Coasters” by Funny Twins Games, 2016).

### 3. Design and Procedure

At the beginning of the experiment, participants completed two arousal-related questionnaires: (1) the ‘Trait’ subscale of the ‘State-Trait Anxiety Inventory’ (STAI-T; Spielberger, 1970; Spielberger, 1989) and (2) the ‘Sensation Seeking’ subscale of the ‘UPPS Impulsive Behaviour Scale’ (UPPS; Whiteside & Lynam, 2001; Schmidt et al., 2008). Participants then had a 280-s VR experience, which included two rollercoasters (153 s, 97 s) and a 30-s intermediate break, twice: once keeping their head straight to avoid movement-related artifacts in the EEG data (*nomov* condition) and once freely moving their head (*mov* condition). Since a high degree of freedom in participants’ movements was expected, the *nomov* condition was introduced to test movement-related noise or artifacts. The order was randomized across participants (in the 29 analyzed, *nomov*- *mov*: *n* = 13; *mov*-*nomov*: *n* = 16). The two rollercoaster rides were chosen for their length, to avoid physical discomfort from wearing the HMD for an extended period, and for their content, to maximize variance in emotional arousal. The first ‘Space’ rollercoaster was a ride in space featuring planets, asteroids and spaceships. The main events were two vertical spins starting around 47 and 73 s after the start of the rollercoaster. The environment contained few explosions of asteroids colliding. Apart from the sounds of explosions, only minimal sound effects were present during the space experience. The second ‘Andes’ rollercoaster was a ride featuring mountain scenery, which included a steep drop (24 s after start), two jumps with steep landings (31 and 67 s after start), two passages through fires under the tracks (20 and 55 s after start) and a looping (60 s after start). A jingling melody played in the background, and sound effects related to fire, airflow and the waggon on the track were also present. In the subsequent rating phase (following immediately after the VR experience for each movement condition), the participants saw a 2D recording of their experience on a virtual screen. While viewing the video, the participants recalled their emotional arousal and continuously reported it using a dial (Griffin PowerMate USB; sampling frequency: 50 Hz), with which they manipulated a vertical rating bar next to the video, ranging from low (0) to high (50) (steps of 1; McCall et al., 2015). No visible notches were labeled on the dial and the visual rating bar (see Figure S1 in Supplements). The participants were instructed as follow: ‘When we show you the video, please state continuously how emotionally arousing or exciting the particular moment during the VR experience was’ (in German: ‘Wenn wir dir das Video zeigen, gebe bitte durchgehend an, wie emotional erregend, bzw. aufregend der jeweilige Moment während der VR Erfahrung war’). After the experiment, the participants were also asked to rate the presence (7-points Likert scale, from 1 to 7) and valence (7-points Likert scale, from -3 to 3) for each rollercoaster in each head movement condition (i.e., four ratings in total per variable). For further details of the experimental setup and data acquisition procedures, please refer to Hofmann, Klotzsche, Mariola et al. (2021).

### 4. Preprocessing

An overview of the preprocessing steps is presented in Figure 1. The preprocessing steps were applied separately for data recorded during the *nomov* and *mov* conditions (i.e., without and with head movement, respectively).

**Figure 1.**
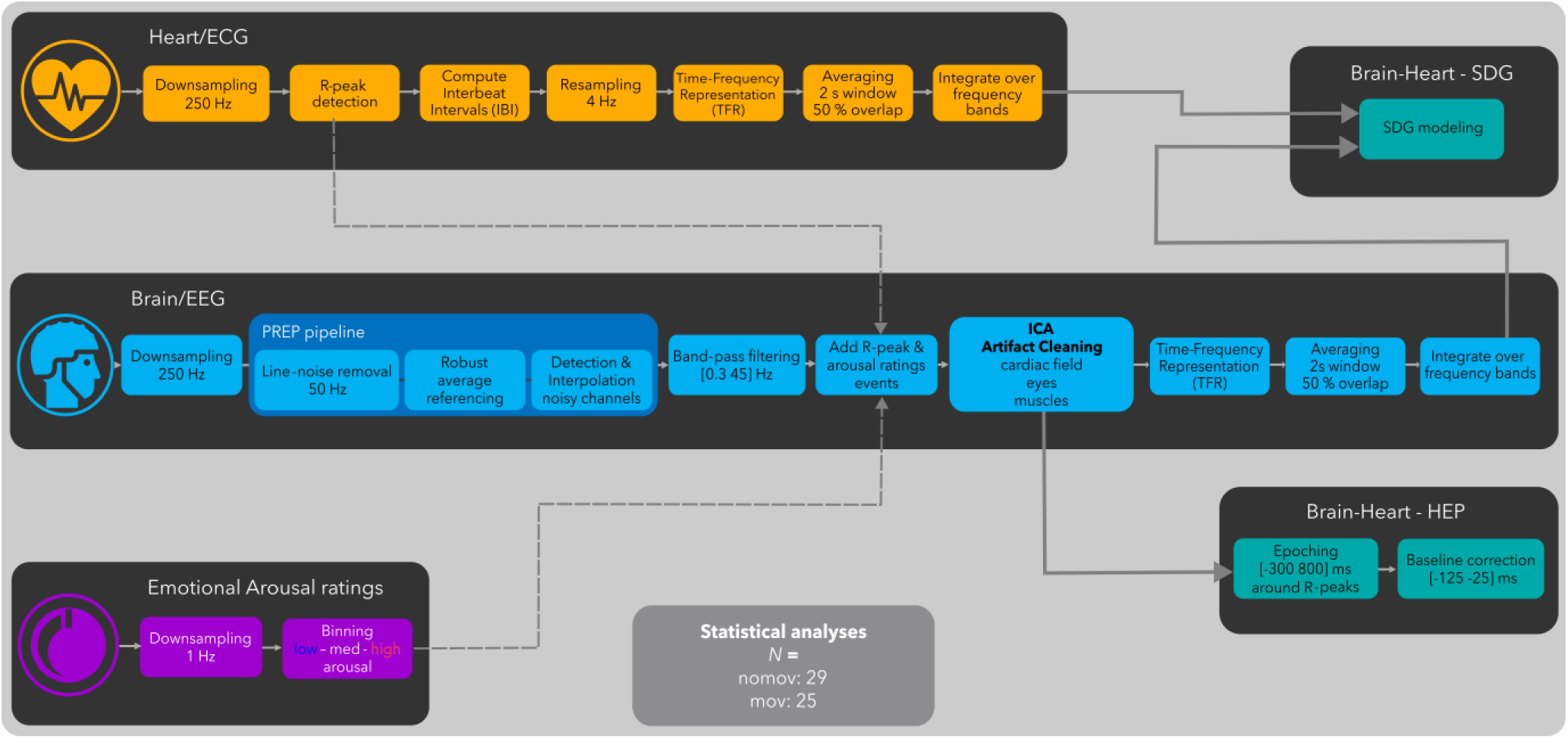
Flowchart of the preprocessing pipeline. Emotional arousal ratings (in purple), Heart/ECG (in orange), Brain/EEG (in blue), Brain-Heart (in turquoise): Heart-evoked potentials (HEP) and synthetic data generation (SDG) modeling; Please note that TFR computation and SDG modeling was preceded by symmetric padding. Further details of the preprocessing steps can be found in the corresponding sections.

#### 4.1. Data Cropping

EEG data, ECG data, and retrospective arousal ratings were cropped by 10 s (2.5 s at beginning and 2.5 s at end of each rollercoaster) to avoid outliers related to the onset and offset of the virtual rollercoaster rides. This resulted in two time series (*nomov, mov*) of 270 s per participant.

#### 4.2. Emotional Arousal Ratings

Subjective reports were downsampled to 1 Hz by averaging non-overlapping sliding windows and then divided by a tertile split into three distinct bins of arousal ratings (low, medium, high) per participant. To increase the sensitivity of our analyses as well as to increase the comparability of our findings with previous literature (e.g., Luft & Bhattacharya, 2015) and our own previous analyses (Hofmann, Klotzsche, Mariola et al., 2021), the medium arousal ratings were discarded. Only the low and high bins were considered, resulting in 180 samples per subject (90 for low, 90 for high emotional arousal).

#### 4.3. Heart/ECG

ECG recordings were downsampled to 250 Hz. Automatic R-peak detection and manual inspection/correction was performed with the EEGLAB extension HEPLAB (version 1.0.1; Perakakis, 2019) in MATLAB (version R2022a).

IBI time series were computed, resampled to 4 Hz (cubic-spline interpolation) and their time-frequency representation (TFR) performed using Continuous Wavelet Transform (CWT, mother wavelet: Morlet, ω0 = 6), using the neurokit2 package (version 0.2.3; Makowski et al., 2021) in Python (version 3.10). To this end, to minimize artifacts due to semi-continuity (i.e., transition effects at the beginning and end of the break), the first R-peak after the beginning and end of the break were removed before resampling. To minimize edge artifacts at the beginning and end of the time-series, a symmetric padding (70 s of inverted data, concatenated at the beginning and end of time-series) was added to the IBI time-series before computing CWT.

The TFRs were downsampled to 1 Hz by averaging within a sliding window of 2 s and 50 % overlap. High and low heart rate variability spectral-power time series were derived by integrating TFRs within the frequency ranges 0.04–0.15 Hz (LF-HRV) and 0.15–0.4 Hz (HF-HRV). Finally, the symmetric padding was partially removed from the IBI and HRV time-series, keeping 35 s of symmetric padding in order to initialize the SDG models (see below).

#### 4.4. Brain/EEG

EEG data were preprocessed and analyzed with custom MATLAB and Python scripts building on the EEGLAB toolbox (version 2023.0; Delorme & Makeig, 2004) and MNE (version 1.1.0; Gramfort et al., 2013; Larson et al., 2022). Continuous data were downsampled to 250 Hz and PREP pipeline (v.0.56.0; Bigdely-Shamlo et al., 2015) procedures were applied for line-noise removal (line frequency: 50 Hz), robust referencing to average, and detection as well as spherical interpolation of noisy channels. On average, 2.08 and 2.47 channels per subject were interpolated in the *nomov* and *mov* condition, respectively. Data were then bandpass filtered (0.3–45 Hz; Hamming windowed sinc FIR filter). Retrospective arousal ratings and R-peak timings were added as event markers to the data sets.

ICA (Extended infomax; Lee et al., 1999) decomposition was used to identify and remove EEG artifacts caused by eye movements, blinks, cardiac field artifacts (CFA) and muscular activity. To facilitate the decomposition, ICA projection matrices were calculated on a copy of the data, high-pass filtered at 1 Hz (instead of 0.3 Hz; Winkler et al., 2015) and from which the noisiest parts had been removed. To this end, a copy of the continuous data was split into 270 epochs of 1 s length. Epochs containing absolute voltage values > 100 µV in at least one channel (excluding channels that reflected eye movements, i.e., EOG channels, Fp1, Fp2, F7, F8) were deleted. Extended infomax (Lee et al., 1999) ICA decomposition was calculated on the remaining parts of the data (after correcting for rank deficiency with a principal component analysis). Subjects with > 90 to-be-deleted epochs (33 % of the data) were discarded from further analyses (*nomov*: *n* = 5; *mov*: *n* = 9). Artefactual ICA components were semi-automatically selected using the SASICA extension (version 1.3.8; Chaumon et al., 2015) of EEGLAB and visual inspection. On average, 10.51 (*nomov;* eye: 2.97, muscle: 4.79, CFA: 1.21, other: 1.45) and 13.40 (*mov;* eye: 3.36, muscle: 5.84, CFA: 1.01, other: 2.92) components per subject were discarded. The remaining ICA weights were back-projected onto the continuous time-series.

TFRs were calculated using Continuous Wavelet Transform (CWT; mother wavelet: Morlet, number of cycles = 7). To minimize edge artifacts and transition effects, CWT was computed on each section separately (roller coaster 1, break, roller coaster 2) with a symmetric padding and then re-concatenated together.

The resulting TFRs were downsampled to 1 Hz by averaging within a moving window of 2 s and 50 % overlap. The EEG spectral power time-series at each electrode were derived by integrating the TFRs within the five classical frequency bands: delta (δ; 0.3–4 Hz), theta (θ; 4–8 Hz), alpha (α; 8–13 Hz), beta (β; 13–30 Hz) and gamma (γ; 30–45 Hz). Thirty-five seconds of symmetric padding at the beginning of the time-series was kept to initialize the SDG models (see below).

#### 4.5. Brain-Heart

In addition to the preprocessing steps described above, additional processing steps were applied for the joint analysis of heart and brain signals:

##### 4.5.1. Heartbeat-Evoked Potentials (HEP)

In the time domain, the preprocessed EEG data were epoched, from 300 ms before each R-peak to 800 ms after. Epochs were then baseline corrected by subtracting for each epoch the mean voltage in the time-window [-125 -25] ms.

4.5.2. Synthetic Data Generation (SDG)

IBI, LF-HRV, HF-HRV, and alpha spectral power time-series for each subject and head movement condition were used as inputs for the SDG model (Catrambone et al., 2019).

In this approach, the ECG R-peaks are modeled (based on integral pulse frequency modulation; Brennan et al., 2002) as a sequence of Dirac delta functions, which are generated by integrating the activity of two oscillators (one for each HRV frequency band: LF and HF) representing the autonomic regulation of the heart activity. Crucially, the amplitudes of the oscillators depend on an additional brain-to-heart coupling coefficient. This coefficient quantifies the strength of the information flow from a specific EEG frequency band to a specific HRV frequency band. The EEG signal, in turn, is modeled (based on adaptive Markov process amplitude; Al-Nashash et al., 2004) as multiple oscillators (one for each frequency band: delta, theta, alpha, beta, gamma), whose amplitudes depend on an additional heart-to-brain coupling coefficient. This coefficient quantifies the strength of the information flow from a specific HRV frequency range to a specific EEG frequency range. Therefore, both EEG and HRV time-series are mutually dependent, and their interaction is modulated by the introduced coupling coefficients. Finally, by means of inverse modeling, both brain-to-heart and heart-to-brain coupling coefficients can be derived.

As model parameters, a 15 s long time window with a 1 s step was used to estimate the coefficients, and the central frequencies used were *ωs* = 2π · 0.1 *rad/s* (LF band central frequency = 0.1 Hz) and *ωp* = 2π · 0.25 *rad/s* (HF band central frequency = 0.25 Hz). The output of the model was the time courses of the coupling coefficients, in both direction (i.e., brain-to-heart and heart-to- brain), for all combinations of brain oscillations (delta, theta, alpha, beta, and gamma) and HRV components (LF and HF).

Please note that with these parameters, the SDG model needed 30 s to initialize. To avoid missing data for the first 30 s of the VR experience, all the inputs had 35 s of symmetric padding added at the beginning and end. This symmetric padding was kept from the previous preprocessing steps (TFR computation).

### 5. Statistical Analysis

#### 5.1. Selection of Region of Interest (ROI)

Based on previous findings (Hofmann, Klotzsche, Mariola et al., 2021; Candia-Rivera et al., 2022), we focused on the interaction between alpha power and HF-HRV, in all parieto-occipital electrodes. We therefore defined a ROI including the following EEG electrodes: Pz, P3, P4, P7, P8, O1, O2, and Oz.

#### 5.2. Heart/ECG

From the IBI and HRV time-series, time-points corresponding to low and high arousal ratings were selected and entered into linear mixed models (LMM; lme4 [version 1.1-29] and lmerTest [version 3.1-3] packages in R [version 4.1.0]; Bates et al., 2014; Kuznetsova et al., 2017), with two factors: (emotional) arousal (two levels: high, low); and (head) movement (two levels: nomov, mov). HRV values were log-transformed in order to approximate a normal distribution.

Each factor and their interaction were entered both as fixed and random effect (i.e., full model; Barr et al., 2013), as follows:

IBI or log(HRV) ∼ 1 + arousal + movement + arousal * movement + (1 + arousal + movement + arousal * movement | ID)

In the fixed effects, an intercept and slopes for the two factors and their interaction were estimated. In the random effects, an intercept and slopes for the two factors and their interaction were estimated within participants.

*P*-values for *F-* and *t*-tests were calculated using the lmerTest ANOVA (type 3) function using Satterthwaite’s method. Estimated marginal means (with 95% confidence intervals) and post hoc pairwise comparisons (with Tukey correction for *p*-values) were computed using the emmeans package (version 1.7.3; Lenth, n.d.).

#### 5.3. Brain/EEG

For each participant and each head movement condition, alpha spectral power was averaged over the electrode ROI. From the alpha spectral power time-series, time-points corresponding to low and high arousal ratings were selected and entered into the same LMM as for the Heart/ECG analysis, with two factors: (emotional) arousal (two levels: high, low); and (head) movement (two levels: nomov, mov). Alpha power values were log-transformed in order to approximate a normal distribution.

Each factor and their interaction were entered both as fixed and random effect, as follows: log(α) ∼ 1 + arousal + movement + arousal * movement + (1 + arousal + movement + arousal * movement | ID)

#### 5.4. Brain-Heart

##### 5.4.1. HEP

In a whole-head analysis, non-parametric cluster-based permutation t-tests were used to compare HEP amplitudes between HA and LA between 250 ms and 450 ms after R-peaks, by pooling the data from both head movement conditions. Previous studies (e.g., Schandry et al., 1986; Al et al., 2020) typically reported HEPs between 250 and 400 ms; we extended this time-window to 450 ms to include Luft and Battacharya (2015) findings. A cluster threshold of *p* = .05 (for clustering data points temporally and spatially adjacent) and 10,000 random permutations (to create the null distribution) were used. Clusters with *p* < .05 (two-tailed) in the permutation test were considered significant.

##### 5.4.2. SDG

For each participant and each head movement condition, heart-to-brain and brain-to-heart couplings (model outputs HF-HRV → α and α → HF-HRV, respectively) were averaged over the electrode ROI. Time-points corresponding to low and high arousal ratings were selected and entered into the same LMM as for the Heart/EEG and Brain/EEG analyses, with two factors: (emotional) arousal (two levels: high, low); and (head) movement (two levels: nomov, mov). Coupling values were log-transformed in order to approximate a normal distribution.

Each factor and their interaction were entered both as fixed and random effect, as follows: log(HF-HRV → α) or log(α → HF-HRV) ∼ 1+ arousal + movement + arousal * movement + (1+ arousal + movement + arousal * movement | ID)

### 6. HEP Source Localization

Exact low-resolution tomography analysis (eLORETA, RRID:SCR_007077; Pascual-Marqui, 2007) was used to localize the sources corresponding to HEP differences between HA and LA. Our pipeline was based on the work of Idaji et al., 2020, who customized the eLORETA implementation of the M/EEG Toolbox of Hamburg (https://www.nitrc.org/projects/meth/). Our forward model was constructed via the New York Head model (Haufe et al., 2014; Huang et al., 2016; Haufe & Ewald, 2019) with approximately 2000 voxels and by using 28 out of 30 scalp electrodes (TP9 and TP10 were removed because they are not contained in the model). Crucially, we constrained our sources to be perpendicular to the cortical surface. Individual HA vs. LA scalp activations were taken as the averaged topography of the difference of HEPs between HA and LA within the time-window of observed emotional arousal-related HEP differences at the group level (328 to 360 ms after R-peak, based on the cluster-based permutation testing). Inverse modeling was computed separately per participant and the L2-normalized source activations were then averaged across all subjects.

### 7. Control and Exploratory Analyses

To test the robustness of our methods and the specificity of our results, we conducted several control analyses.

#### 7.1. ECG Waveform

To control for potential confounds in our HEP results, we tested for possible differences in the ECG waveforms between HA and LA. We compared the ECG signal time-locked to the R-peak of HA vs. LA within participants, using two-tailed paired t-tests at all time-points within the time-window of observed emotional arousal-related HEP differences. The *p*-value was corrected for multiple comparisons with False Discovery Rate (FDR; Benjamini & Yekutieli, 2001).

#### 7.2. Other frequency bands

To evaluate the specificity of the effects in the alpha band and explore arousal-related changes in (A) brain activity and (B) BHI (Candia-Rivera et al., 2022), we performed a whole-scalp analysis for (A) all the EEG frequency bands: delta (δ; 0.3–4 Hz), theta (θ; 4–8 Hz), alpha (α; 8–13 Hz), beta (β; 13–30 Hz) and gamma (γ; 30–45 Hz), and (B) their integration in SDG (LF-HRV → δ, δ → LF-HRV, HF-HRV → δ, δ → HF-HRV; LF-HRV → θ, θ → LF-HRV, HF-HRV → θ, θ → HF-HRV; LF-HRV → α, α → LF-HRV, HF-HRV → α, α → HF-HRV; LF-HRV → β, β → LF-HRV, HF-HRV → β, β → HF-HRV; LF-HRV → γ, γ → LF-HRV, HF-HRV → γ, γ → HF-HRV). To this end, at each electrode, we averaged the different metrics over the head movement conditions, and performed paired t-tests between the mean during HA and the mean during LA for each participant.

## Results

### 1. Participants

After the EEG quality assurance during preprocessing, the data from 29 participants (16 men, mean age: 26±3, range: 20-31 years) entered the subsequent analyses. More specifically, after excluding 9 (*mov*) and 5 (*nomov*) participants, results were based on: *Nmov = 25* and *Nnomov = 29*.

### 2. State and trait reports

The Trait Anxiety scores (STAI-T, ‘Trait’ subscale, range of possible scores 20-80) averaged across all participants were 44.34 ± 3.56 (M ± SD, range: 37-51) and the Sensation Seeking scores (UPPS, ‘Sensation Seeking’ subscale, range of possible scores: 1-4) were 3.01 ± 0.38 (M ± SD, range: 2.08-3.67).

The presence ratings averaged across all participants were 4.53 ± 1.81 (M ± SD, range: 1-7) in the *mov* and 4.34 ± 1.80 (M ± SD, range: 1-7) in the *nomov* condition. There were no significant differences between the two conditions (paired t-test; *t*(57) = 1.2, *p* = .225).

The valence ratings averaged across all participants were 1.36 ± 1.42 (M ± SD, range: -3-3) in the *mov* and 1.16 ± 1.52 (M ± SD, range: -3-3) in the *nomov* condition. There were no significant differences between the two conditions (paired t-test; *t*(57) = 1.4, *p* = .176).

### 3. Emotional arousal ratings

The emotional arousal ratings averaged across all timepoints and participants were 27 ± 12 (M ± SD, range: 0-50) in the mov and 24 ± 13 (M ± SD, range: 0-50) in the nomov condition. Descriptively, the emotional arousal was highest for the Andes Coaster, lower for the Space Coaster, and lowest for the break (see Figure 5 [z-scored] and S2).

### 4. Heart/ECG

There was a significant main effect of arousal (*F*(1, 28.6) = 5.9, *p* = .021) on HF-HRV (see Figure 2). No evidence for a main effect of head movement (*F*(1, 24.6) = 1.6, *p* = .218), nor the interaction (F(28.1) = 1.4, *p* = .245) was found. Post-hoc pairwise comparisons of the estimated marginal means revealed significantly lower HF-HRV for HA compared to LA in the free head movement condition (*mov*; *t*(24.7) = -3.0, *p* = .007), but not in the condition without head movement (*nomov*; *t*(28.0) = -1.1, *p* = .285).

**Figure 2.**
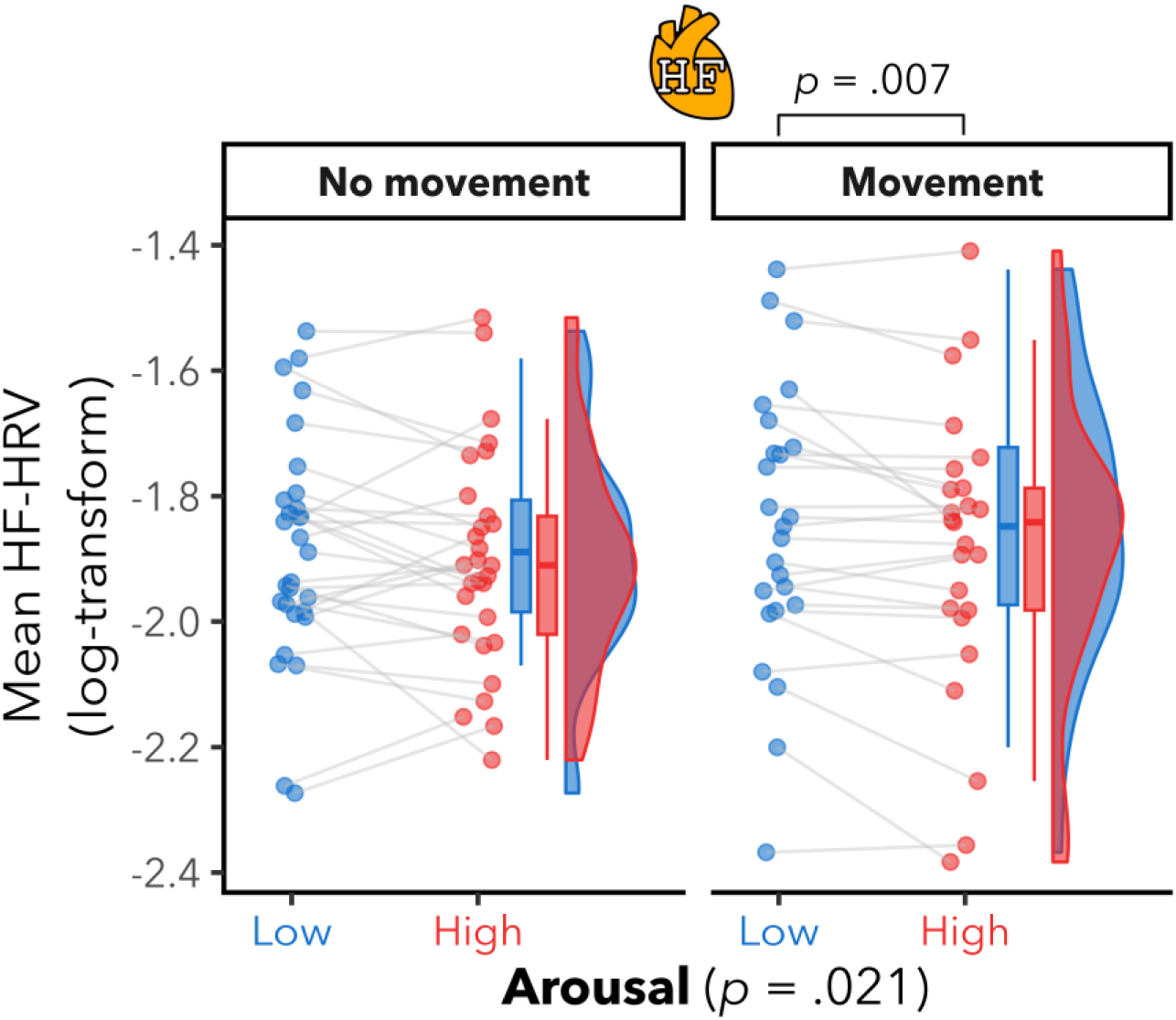
Significant high-frequency heart rate variability (HF-HRV) differences between high and low emotional arousal, in the free head movement condition. Mean HF-HRV per participant for low (blue color) and high (red color) arousal and for runs without and with free head movement. Box plots (horizontal bar: mean; whiskers: 1.5 interquartile range); individual dots represent individual participants. There was a significant effect of arousal, but not of head movement nor the interaction. Post-hoc pairwise comparisons showed lower HF-HRV during higher arousal in the free head movement condition, but not in the without head movement condition. There was no evidence for any significant effects of arousal, movement nor interaction on interbeat intervals (IBI) (see Figure S3 in Supplements). There was also a significant effect of arousal on LF-HRV (see Figure S3 in Supplements), but no significant effects of movement nor interaction.

We observed the same patterns for LF-HRV, that is, a significant main effect of arousal *F*(1, 26.9) = 9.5, *p* = .005; see Figure S3 in Supplements), no significant main effect of movement (*F*(1, 25.5) = 2.2, *p* = .152), and no significant arousal-by-movement interaction (*F*(1, 26.0) = 0.5, *p* = .492). Post-hoc pairwise comparisons of the estimated marginal means revealed significantly lower LF-HRV during higher arousal in the *mov* (*t*(25.1) = -3.2, *p* = .004), but not in the *nomov* (*t*(28.0) = -1.6, *p* = .124) condition. No significant effects were observed on heart rate (i.e., IBI; all *p* > .05; see Figure S3 in Supplements).

### 5. Brain/EEG

We found a significant effect of arousal (*F*(1, 27.7) = 30.7, p < .001) on participants’ alpha power in parieto-occipital regions (see Figure 3). There were no significant head movement (*F*(1, 24.1) = 2.5, *p* = .128) nor interaction effects (*F*(1, 24.1) = 2.9, *p* = .101). Post-hoc pairwise comparisons of the estimated marginal means revealed significantly lower alpha power in parieto-occipital regions for HA compared to LA in both the free (*mov*; *t*(25.9) = -3.2, *p* = .004) and without (*nomov*; *t*(28.0) = -6.6, *p* < .001) head movement conditions.

**Figure 3.**
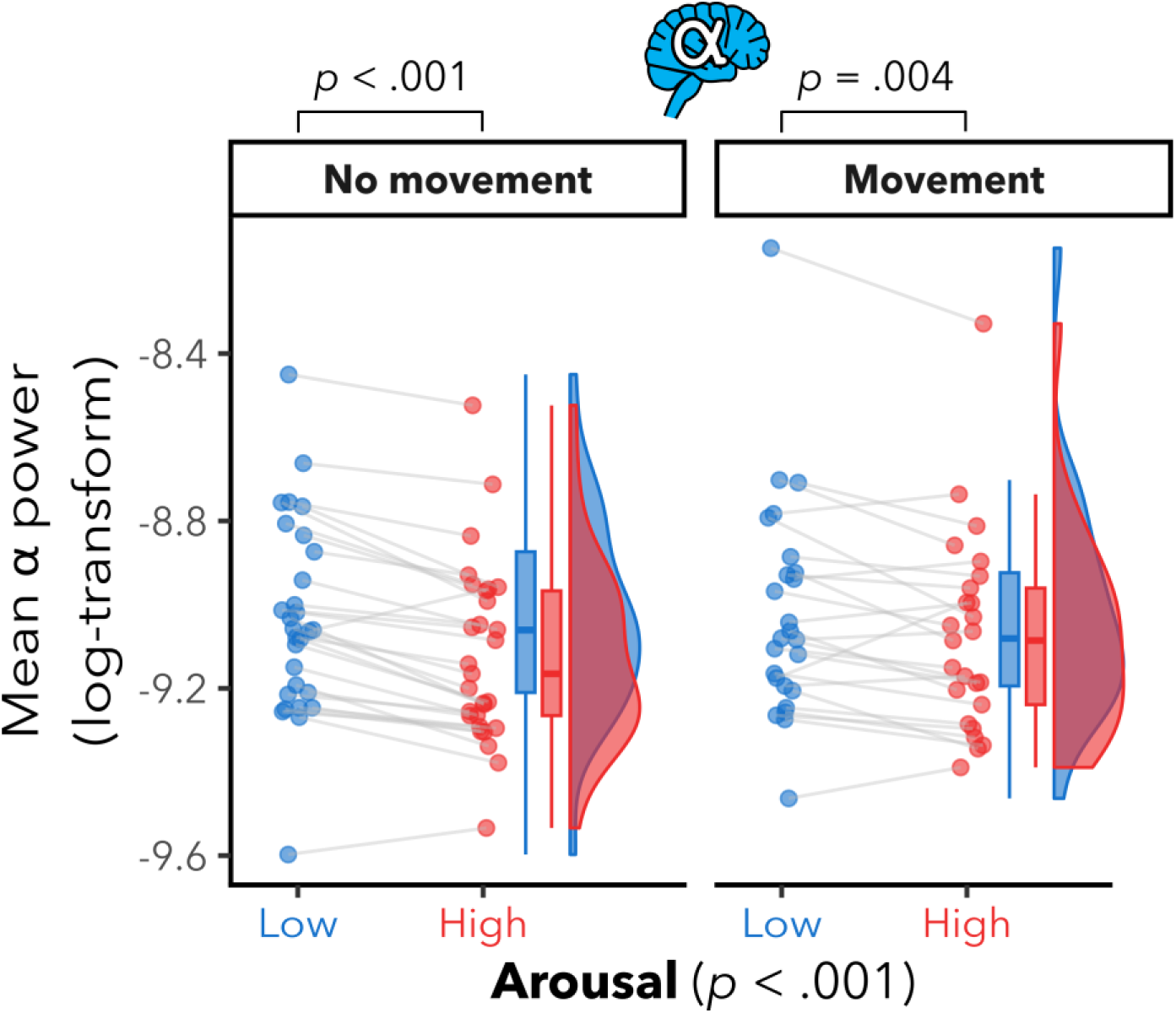
Lower parieto-occipital alpha power in high vs. low arousal. Mean log α power within the region of interest (ROI: electrodes Pz, P3, P4, P7, P8, O1, O2, and Oz) per participant for low (blue color) and high (red color) arousal and for runs with and without free head movement. Box plots (horizontal bar: mean; whiskers: 1.5 interquartile range); individual dots represent individual participants. There were significant effects of arousal, but not of head movement nor interaction. Post-hoc pairwise comparisons showed significantly lower alpha power in parieto-occipital regions for HA compared to LA in both the free and without head movement conditions.

### 6. Brain-Heart

#### 6.1. HEP

The cluster-based permutation tests revealed a significant HEP difference between HA and LA, indicated by a cluster at the left fronto-central regions (C3, FC1, FC5, Fz, F3 electrodes; Monte Carlo *p* = .019; see Figure 4A) from 328 to 360 ms after R-peak, with lower (i.e., more negative) HEP amplitude for HA than for LA (see Figure 4B).

**Figure 4.**
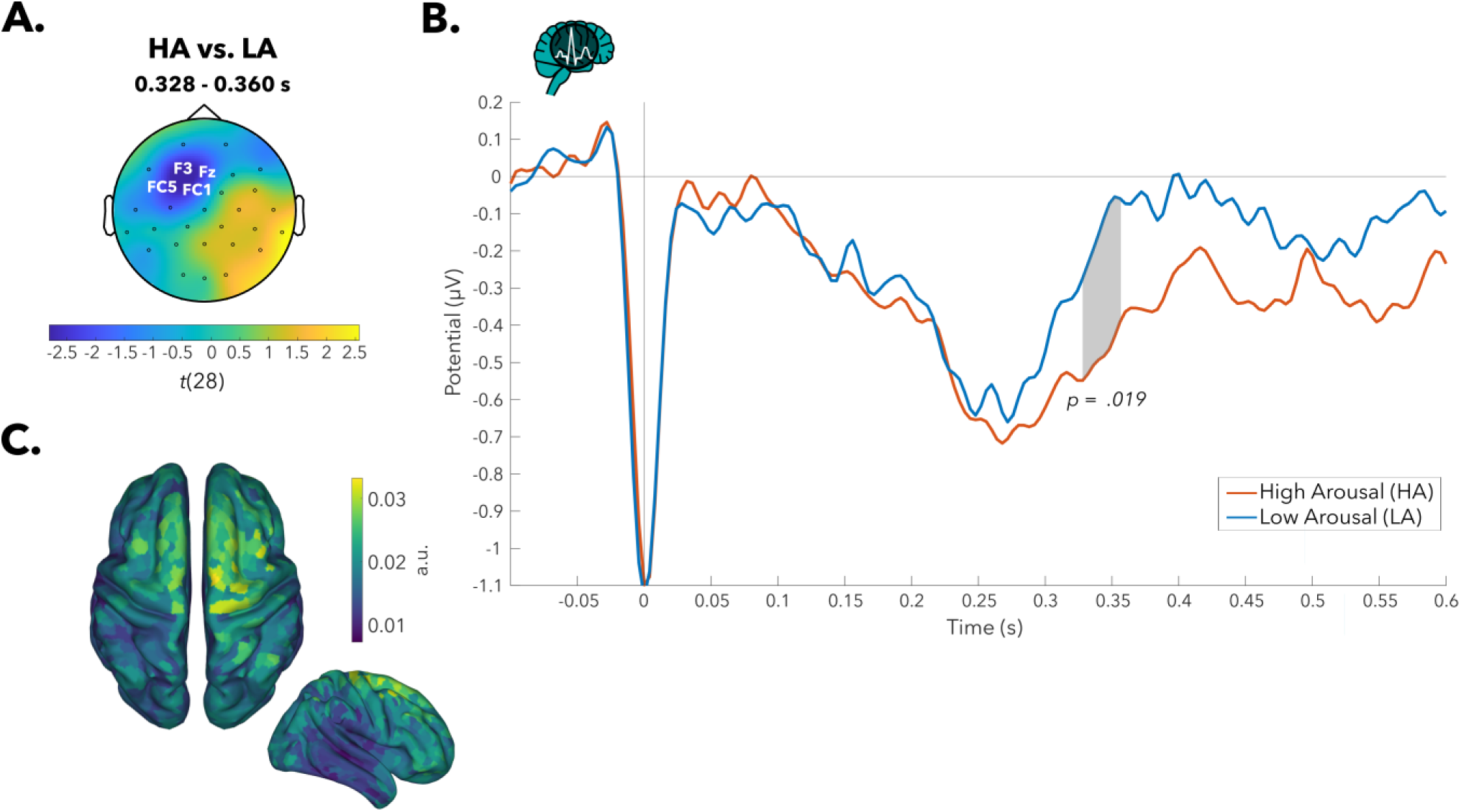
Heartbeat-evoked potential (HEP) amplitudes significantly differed between high (HA) and low arousal (LA) over left fronto-central electrodes. **A.** Topographical map of t-values for HEP differences between HA and LA: Grand average across 29 participants (pooled data across both head movement conditions) in the 328- to 360-ms time window, where a significant difference (HA > LA) was observed in the cluster of highlighted electrodes. **B.** HEP time courses (HA in red, LA in blue) averaged across the cluster. **C.** Source localization (exact low resolution tomography analysis [eLORETA]) of HEP differences between HA and LA. The projection in source space suggests a distribution of sources with strongest values in premotor, sensorimotor, supplementary motor areas, around the central sulcus. Colors represent the inversely modeled contribution of the cortical voxels to the spatial pattern yielded by the HA vs. LA contrast.

The source localization (via eLORETA) yielded a distribution of sources where the strongest values were located close to and inside the central sulcus, in premotor, sensorimotor and supplementary motor areas (see Figure 4C).

We also observed another cluster over right temporo-parietal electrodes (P8, TP10, T8 electrodes, from 352 to 356 ms after R-peak), with higher HEP amplitudes for HA compared to LA. This cluster, however, did not survive cluster-correction for multiple comparisons (Monte Carlo *p* = .310).

#### 6.2. SDG

For an overview, the mean time-series over all participants of heart, brain and brain-heart metrics of interest, for the condition with free head movement (*mov*), are presented in Figure 5. The time-series for the condition without head movement (*nomov*) are available in the Supplements (see Figure S4).

**Figure 5.**
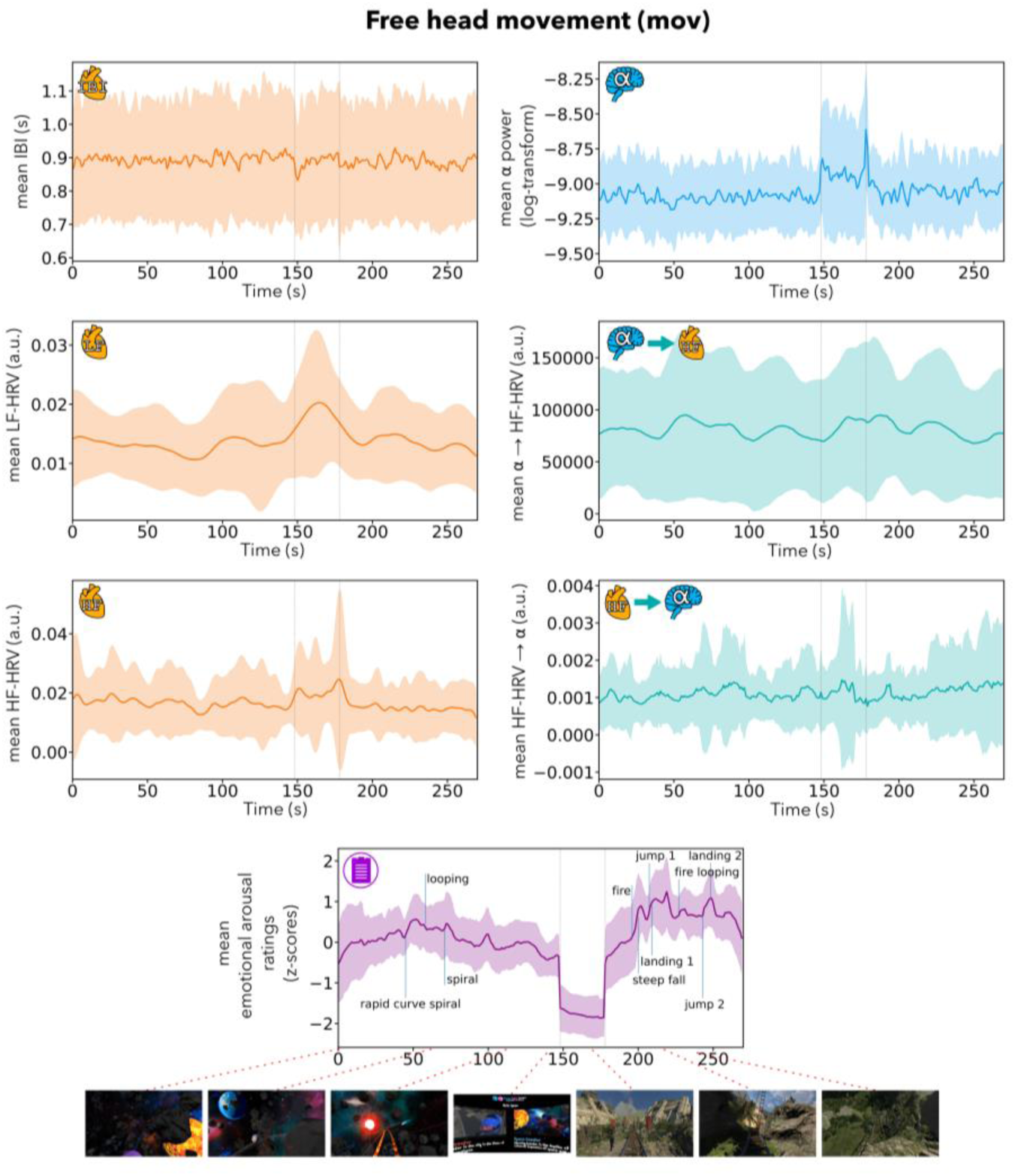
Mean time-series across participants of interbeat intervals (IBI), low-frequency heart rate variability (LF-HRV), high-frequency heart rate variability (HF-HRV), log α power averaged across the region of interest (ROI: electrodes Pz, P3, P4, P7, P8, O1, O2, and Oz), brain-to-heart coupling coefficient (α → HF-HRV; averaged across ROI), heart-to-brain coupling coefficient (HF-HRV → α; averaged across ROI) and emotional arousal ratings, for the condition with free head movement (mov). The shaded areas represent +/− 1 SD. Coloured lines: mean across participants; vertical lines (light grey): beginning and end of the break; vertical lines (blue): manually labeled salient events (for illustration). Bottom row: exemplary screenshots of the virtual reality (VR) experience. The time-series for the condition without head movement (nomov) are shown in the Supplements (see Figure S4).

There was no significant main effect of arousal (HF-HRV → α: *F*(1, 27.1) = 0.01, *p* = .934; α → HF-HRV: *F*(1, 22.3) = 0.3, *p* = .619) on the BHI coupling coefficients in both directions (see Figure 6). There was also no evidence for a main effect of movement (HF-HRV → α: *F*(1, 24.3) = 0.8, *p* = .371; α → HF-HRV: *F*(1, 23.5) = 1.0, *p* = .339) nor an arousal-by-movement interaction (HF-HRV → α: *F*(1, 25.2) = 1.6, *p* = .217; α → HF-HRV: *F*(1, 22.7) = 0.01, *p* = .930).

**Figure 6.**
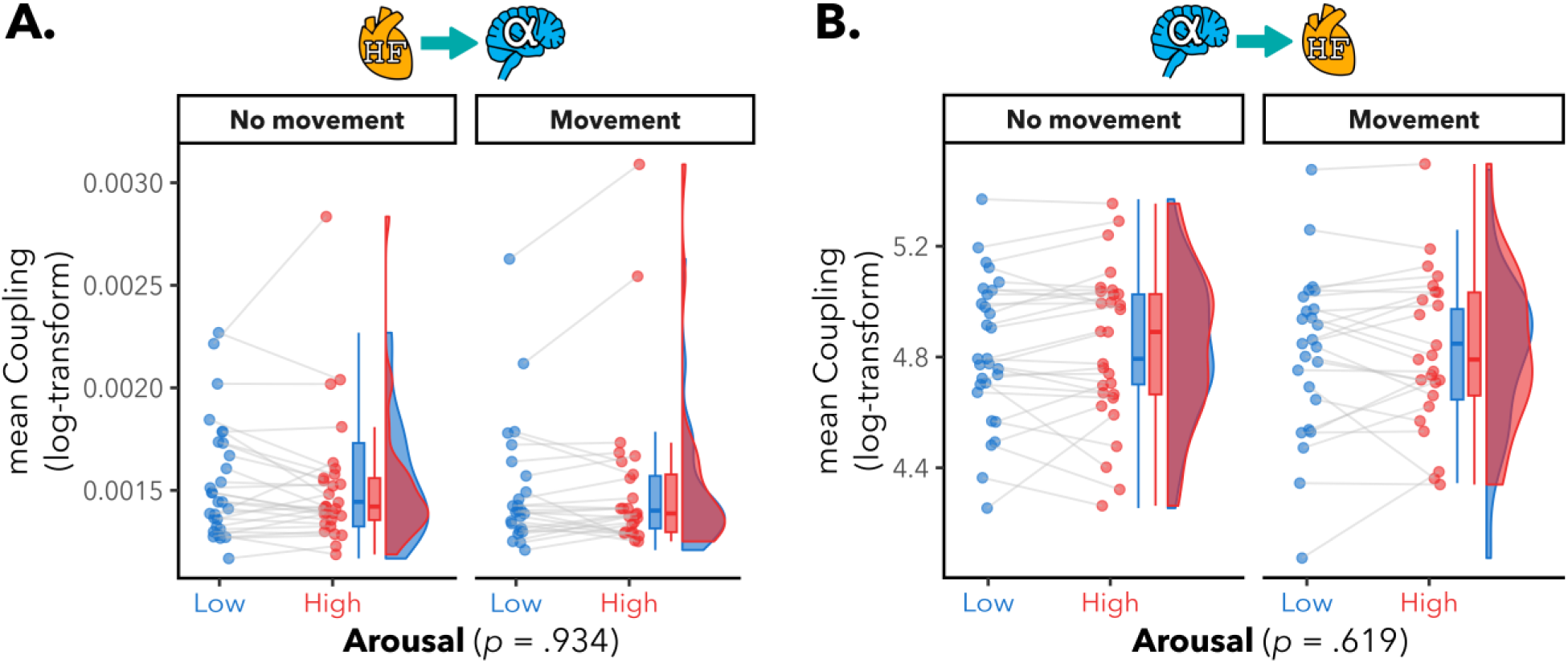
No significant Brain-Heart Interaction (BHI) differences between high and low emotional arousal. Mean directional BHI coupling coefficients per participant, for low (blue color) and high (red color) arousal, and for runs with and without free head movement. The BHI coupling coefficients were computed using a Synthetic Data Generation (SDG) model (Catrambone et al., 2019); Heart related input: high-frequency heart rate variability (HF-HRV); Brain related input: α power within ROI. Box plots (horizontal bar: mean; whiskers: 1.5 interquartile range); individual dots represent individual participants. **A.** Heart-to-brain coupling coefficient (HF-HRV → α; averaged across ROI). There was no evidence for any significant effects of arousal, movement nor interaction on HF-HRV → α (all p > .05). **B.** Brain-to-heart coupling coefficient (α → HF-HRV; averaged across ROI) during HA than LA. There was no evidence for any significant effects of arousal, movement nor interaction on α → HF-HRV (all p > .05).

### 7. Control and Exploratory Analyses

#### 7.1. ECG waveform

There were no significant differences (max *t*(28) = 1.1, min *p* = .27, FDR-corrected) in the ECG waveform between HA and LA conditions at all the time-points within the time window of EA related HEP differences (328 to 360 ms after R-peak).

#### 7.2. Other frequency bands

A summary of the results can be found in Table 1 and figures including topographies in the Supplements.

**Table 1.**
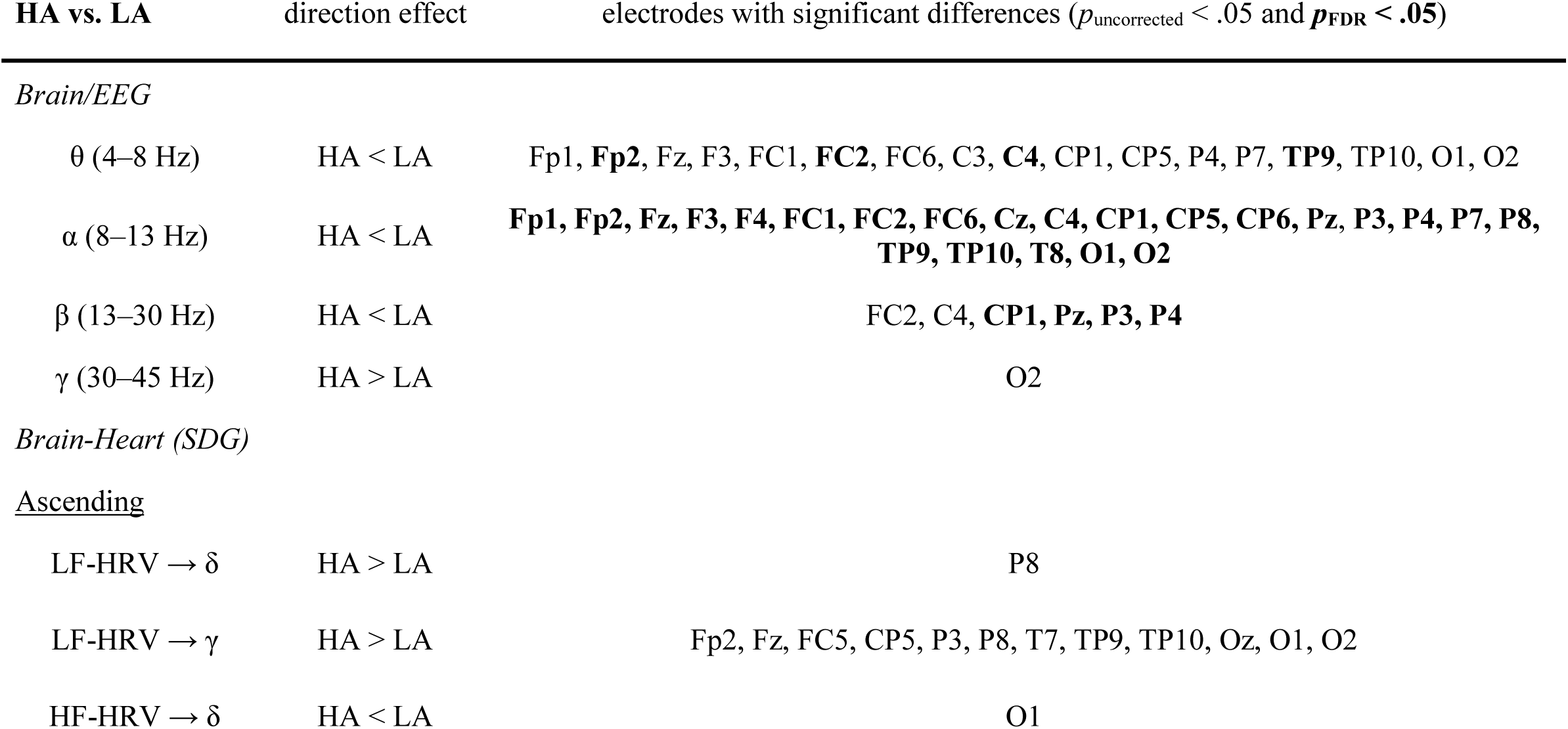
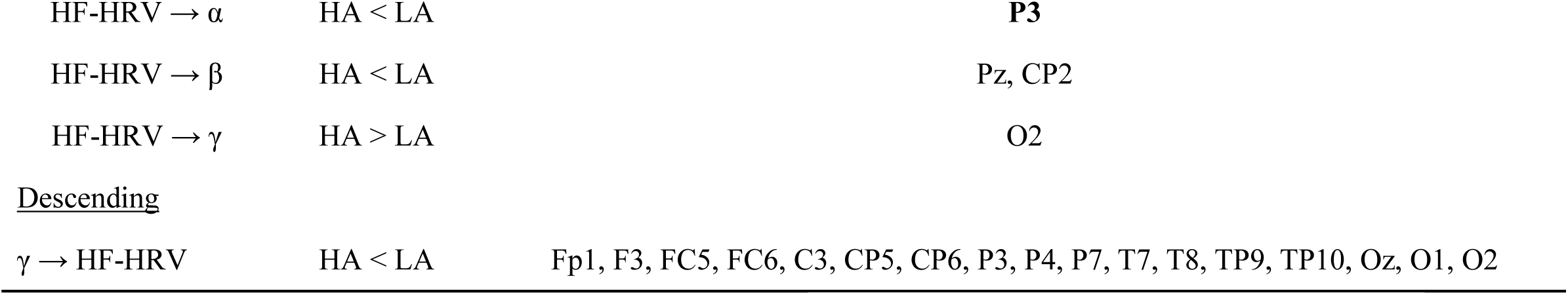
Summary of the exploratory analyses. At each electrode, the different metrics were averaged over the head movement conditions, and paired t-tests were performed between the mean during high arousal (HA) and the mean during low arousal (LA) for each participant. Topographies can be found in the Supplements (see Figure S5 and S6). SDG: synthetic data generation modeling; LF/HF-HRV: low/high-frequency heart rate variability.

| HA vs. LA | direction effect | electrodes with significant differences ($p_{\text{uncorrected}} < .05$ and $p_{\text{FDR}} < .05$) |
| --- | --- | --- |
| *Brain/EEG* |  |  |
| $\theta$ (4–8 Hz) | HA < LA | Fp1, **Fp2**, Fz, F3, FC1, **FC2**, FC6, C3, **C4**, CP1, CP5, P4, P7, **TP9**, TP10, O1, O2 |
| $\alpha$ (8–13 Hz) | HA < LA | **Fp1, Fp2, Fz, F3, F4, FC1, FC2, FC6, Cz, C4, CP1, CP5, CP6, Pz, P3, P4, P7, P8, TP9, TP10, T8, O1, O2** |
| $\beta$ (13–30 Hz) | HA < LA | FC2, C4, **CP1, Pz, P3, P4** |
| $\gamma$ (30–45 Hz) | HA > LA | O2 |
| *Brain-Heart (SDG)* |  |  |
| <u>Ascending</u> |  |  |
| LF-HRV $\rightarrow \delta$ | HA > LA | P8 |
| LF-HRV $\rightarrow \gamma$ | HA > LA | Fp2, Fz, FC5, CP5, P3, P8, T7, TP9, TP10, Oz, O1, O2 |
| HF-HRV $\rightarrow \delta$ | HA < LA | O1 |
| HF-HRV $\rightarrow \alpha$ | HA < LA | **P3** |
| HF-HRV $\rightarrow \beta$ | HA < LA | Pz, CP2 |
| HF-HRV $\rightarrow \gamma$ | HA > LA | O2 |
| Descending |  |  |
| $\gamma \rightarrow$ HF-HRV | HA < LA | Fp1, F3, FC5, FC6, C3, CP5, CP6, P3, P4, P7, T7, T8, TP9, TP10, Oz, O1, O2 |

Exploring arousal-related changes in whole-brain activity, we found different patterns of activation for HA vs. LA (see Figure S5 in Supplements). For the theta band, we found 17 EEG electrodes (see Table 1) with lower theta power during higher arousal. For the alpha band, there were 23 EEG electrodes with lower alpha power during higher arousal. For the beta band, there were 6 EEG electrodes with lower beta power during higher arousal. For the gamma band, one EEG electrode showed higher gamma power during higher arousal.

Exploring further arousal-related changes in BHI, we found different patterns of activation for HA vs. LA (see Figure S6 in Supplements). For HF-HRV → δ, one EEG electrode (see Table 1) showed lower BHI coupling coefficients during higher arousal. For LF-HRV → δ, one EEG electrode showed higher BHI coupling coefficients during higher arousal. For HF-HRV → α, one EEG electrode showed lower BHI coupling coefficients during higher arousal. For HF-HRV → β, two EEG electrodes showed lower BHI coupling coefficients during higher arousal. For HF-HRV → γ, one EEG electrode showed higher BHI coupling coefficients during higher arousal. For LF-HRV → γ, there were 12 EEG electrodes (see Table 1) with higher BHI coupling coefficients during higher arousal. For γ → HF-HRV, there were 17 EEG electrodes with lower BHI coupling coefficients during higher arousal. Furthermore, concerning the gamma band, when looking at the interception (FC5, CP5, P3, T7, TP9, TP10, O1, Oz, O2) of both significant electrodes for the ascending (LF-HRV → γ) and descending (γ → HF-HRV) directions, both effects were still present (LF-HRV → γ: *t*(28) = 2.8; *p* = .008; γ → HF-HRV: *t*(28) = -2.7; *p* = .012).

## Discussion

We investigated brain-heart interactions during an emotionally arousing experience in immersive VR. In particular, we analyzed emotional arousal-related cardiac activity (particularly its vagal regulation) and the extent to which it contributed to the previously reported link between emotional arousal and parieto-occipital alpha power (Hofmann, Klotzsche, Mariola et al., 2021). We found differences in heart activity, brain activity and - less consistently - brain-heart interactions between states of HA and LA. More specifically, we observed significant arousal-related BHI differences in event-related analyses (i.e., HEPs) but not in oscillatory analyses (i.e., SDG modeling), although whole-brain exploratory analyses pointed towards increased ascending heart-to-brain (i.e., LF-HRV → γ) and reduced descending brain-to-heart (i.e., γ → HF-HRV) functional information flow during higher emotional arousal. Generally, our findings extend previous results from classical studies and confirm the link between emotional arousal and HF-HRV, parieto-occipital alpha power, and HEP amplitude under more naturalistic conditions.

### 1. Heart/ECG

Analyzing the heart signal, we found significant effects of emotional arousal on both LF- and HF-HRV. This is in line with previous research showing lower HF-HRV during states of higher emotional arousal (Valenza et al., 2012; Luft & Bhattacharya, 2015; Hildebrandt et al., 2016). As HF-HRV reflects vagal cardioregulation (Task Force, 1996), a possible interpretation is that the parasympathetic regulation of heart activity decreased (i.e., vagal withdrawal) during higher emotional arousal. Fluctuations in HF-HRV during emotion have been associated with functional changes in medial prefrontal cortex (Lane et al., 2009), a brain region shared by the CAN and the SN. The changes in HF-HRV during emotional arousal could therefore be also linked to a (bottom-up) modulation of attention in presence of emotional stimuli.

Concerning LF-HRV, the measure reflects both the sympathetic and parasympathetic influences on the heart rate. While the emotional arousal-related changes in LF-HRV could thus be driven by changes in parasympathetic activity, it remains unclear why we did not find evidence for heart rate differences between LA and HA. As both ANS branches can be active at the same time (Koizumi et al., 1983; Paton et al., 2005), their contribution to the heart rate may have canceled each other out (to only be visible in the variability measures LF- and HF-HRV). Because the heart rate and its sympathetic regulation are known to change more slowly (order of few seconds) than its parasympathetic regulation (order of ms), it is also possible that our analysis (with 1 s resolution) did not capture these delayed changes. Future studies may consider directly assessing the sympathetic regulation of the heart and its potential link to emotional arousal, for instance by modeling regulations at different timescales to extract a sympathetic activity index (Valenza et al., 2018) from the ECG signal and take into account the potential delay of the sympathetic regulation, or measuring pre-ejection periods with impedance cardiography.

### 2. Brain/EEG

Analyzing the brain activity, we confirmed our previous findings of lower parieto-occipital alpha power for HA compared to LA states (Hofmann, Klotzsche, Mariola et al., 2021) with a different statistical approach, using LMM instead of decoding. Of note, exploratory analyses suggest that this effect is not restricted to parieto-occipital cortical areas. This is also in line with previous research using event-related designs (Koelstra et al., 2012; Luft & Bhattacharya, 2015). Because alpha power has also been linked to attentional processes (Klimesch, 2012; van Diepen et al., 2019) and to the encoding of the emotional salience of stimuli (e.g., in orbitofrontal cortex; Todd et al., 2014), the change in alpha power might indicate a (bottom-up) modulation of attention in presence of emotional stimuli (Sutherland & Mather, 2018) - which may be present with a more widespread topography.

As EEG is acquired at the scalp, the posterior alpha rhythm mainly represents activity in parieto-occipital cortices, as shown by EEG-based source reconstruction (e.g., Hofmann, Klotzsche, Mariola et al., 2021) and simultaneous EEG-fMRI (Moosmann et al., 2003; Omata et al., 2013). However, brain regions that are part of the CAN (Benarroch, 1993; Thayer & Lane, 2000, 2009) have also been found to contribute to the posterior alpha rhythm, such as the anterior cingulate cortex and the amygdala (Omata et al., 2013), together with brainstem regions and thalamic nuclei (Omata et al., 2013; Mossmann et al., 2003). Particularly thalamo-cortical loops have been related to the posterior alpha rhythm (Schreckenberger et al., 2004; Hindriks & van Putten, 2013) and this structural overlap between the CAN and the posterior alpha rhythm network may enable (emotional) arousal-related changes in sensory (particularly visual) or attentional processing (Shibata et al., 2024).

In additional exploratory analyses, we investigated emotional arousal-related differences in other frequency bands. We observed significant effects in other frequency bands, such as lower theta power (cf. Aftanas et al., 2002), lower beta power (Schubring et al., 2020; Kim et al., 2021), and higher gamma power (Cao et al., 2020) during HA vs. LA, but these findings should be interpreted with caution given the exploratory nature of these analyses and the risk of false positives.

The main aim of this work was to investigate if these changes in parieto-occipital alpha oscillations and the changes we observed in heart activity (i.e., in HF-HRV) were related to each other. Given the alpha reduction during HA compared to LA, how did vagal cardioregulation come into play?

### 3. Brain-Heart

Combining heart and brain activities together in a multimodal analysis, we investigated BHI during different states of emotional arousal. Using two different approaches, an event-related and an oscillatory analysis, we found - though not consistently across both approaches - that the functional coupling between the heart and the brain was linked to emotional arousal.

Looking at HEPs (event-related analysis), we found a significantly lower (or more negative) HEP amplitude for high compared to low arousal in left fronto-central electrodes. Because there was no significant difference in heart rate (i.e., IBIs) nor in ECG waveform between HA and LA, this HEP difference may not be due to residual CFA nor to a difference of duration between heartbeats. We did not replicate the findings from Luft & Battacharya (2015), in that we did not find a significantly higher HEP amplitude in parieto-occipital regions during higher arousal (a pattern, which we only observed in a non-significant cluster). The topography of the fronto-central cluster, however, might reflect the anterior pole of a dipole that is also visible at the parietal electrodes. Particularly, a separation with an angle of approximately 45° between the positive and negative pole on the scalp is often characteristic of a tangential equivalent dipolar source located inside the central sulcus (e.g., for the somatosensory evoked potential N20; Scherg et al., 2019). Our source localization results support this view by pointing towards a distribution of sources around the central sulcus, in sensorimotor areas. The left fronto-central topography observed here and the right parieto- occipital topography reported by Luft and Battacharya (2015) may thus reflect similar underlying sources.

Our HEP results are also in line with a meta-analysis (Coll et al., 2021) that found a large effect of arousal (including but not restricted to *emotional* arousal) on HEP amplitudes, with the strongest effect around 250 ms after R-peak and in fronto-central electrodes (Cz, C1, C2, C3, C4, FCz, FC1, FC2, FC3, FC4, FC5, FC6 and AFz). More specifically related to *emotional* arousal, a recent study (Marshall et al., 2019) included in this meta-analysis found a HEP suppression for angry vs. neutral faces when they were repeatedly presented. This could reflect a different weighing of exteroceptive vs. interoceptive information to facilitate rapid perceptual processing and behavioral responses, as well as to mobilize (e.g., metabolic) resources (Gianaros & Wager, 2015).

In the confirmatory oscillatory analysis using SDG modeling, we did not find evidence for an association between emotional arousal and directional communication between the brain (indexed by parieto-occipital alpha power) and the heart (indexed by HF-HRV). The results of the SDG whole-scalp exploratory analysis of multiple frequency bands indicated that emotional arousal was associated with specific changes in information flow from the heart to the brain (higher LF-HRV → γ in a temporo-occipital cluster, lower information flow from HF-HRV to the brain in single electrodes) and - less so - from the brain to the heart (lower γ → HF-HRV in a temporo-occipital cluster; Table 1 for details). Although this should, again, be interpreted with caution, these results suggest a link between the level of emotional arousal and the coupling between HRV and brain activity in different frequency bands. Particularly, ascending signals of heart activity seem to inform brain activity and modulate the emotional experience (Candia-Rivera et al., 2022). Our results are partly consistent with the findings of Candia-Rivera and colleagues (2022), who reported higher (ascending) heart-to-brain information flow - although in different frequency bands - during higher arousal, and lower (descending) brain-to-heart information flow during emotion elicitation compared to rest. Another study (Catrambone et al., 2022) using the same model, found that the complexity (operationalized as Fuzzy Entropy) of heart-to-brain information flow was increased during emotional elicitation (via pictures) compared to rest. Interestingly, another recent study (Agrimi et al., 2023) found, along with higher HF-HRV and EEG gamma band power, higher ascending information flow from LF-HRV to gamma in genetically modified mice with chronically elevated (yet within physiological range) heart rate compared to control mice.

Overall, these results extend our previous contribution to the physiology of emotional experience using immersive VR, with our previous “brain-only” results of EEG-derived parieto-occipital alpha power (Hofmann, Klotzsche, Mariola et al., 2021) now complemented by considering the rest of the body (here: the heart or ANS). During states of higher emotional arousal, not only parieto-occipital alpha power was lower but so were LF-HRV, HF-HRV, and HEP amplitudes over fronto-central electrodes. While we did not find evidence for the hypothesized changes in BHI between parieto-occipital alpha power and HF-HRV (in either direction), exploratory analyses suggest several other emotional arousal-related BHI changes, notably in temporo-occipital gamma power, where higher emotional arousal was linked to decreased brain-to-heart (γ → HF-HRV) and increased heart-to-brain (LF-HRV → γ) information flow. Thereby, heart-to-brain information flow seems to change more broadly (in time and space) with different affective states than - in the opposite direction - the brain-to-heart information flow. This supports the view that signals from the internal body (e.g., the heart) influence our perception of the world and our interaction with it (Ohl et al., 2016; Kunzendorf et al., 2019; Motyka et al., 2019; Galvez-Pol et al., 2022). Changes in bodily rhythms, as they occur in different affective states, change attentional processes (Sutherland & Mather, 2018) or - more generally - the way in which sensory evidence is accumulated (Allen et al., 2022), for example through increased ascending (heart-to-brain) compared to reduced descending (brain-to-heart) information flow.

### 4. Limitations and Future Directions

Several limitations should be considered when interpreting our findings. It should be noted that the sample size was relatively small and consisted of young healthy participants. This limits the generalizability of our findings to the whole population. While the dataset is relatively short and potentially noisy due to the more naturalistic conditions of data acquisition, this is a tradeoff that comes with using a more realistic affective stimulation.

Other limitations are due to the approach of a secondary analysis - the experimental design was not optimized for some analysis methods used here. For example, to avoid transition effects between the different parts of the VR experience, data needed to be trimmed at the beginning and end of rollercoasters. This discontinuity was not ideal for time-resolved metrics such as heart rate, HRV and SDG couplings. Fully continuous data without trimming would be preferable in future experiments.

Furthermore, the emotional arousal ratings were binned into HA and LA in order to increase sensitivity and to maximize comparability to previous findings (Luft & Bhattacharya, 2015) as well as our previous analyses (Hofmann, Klotzsche, Mariola et al., 2021). Models that include the continuous ratings (as also part of Hofmann, Klotzsche, Mariola et al., 2021) can provide a more fine-grained picture of the relationship between emotional arousal and physiological measures.

As respiration modulates vagal cardioregulation (Benarroch, 1993) and HEP amplitudes (Zaccaro et al., 2022), we cannot exclude potential emotional arousal-related changes in respiration (e.g., its rate) and assess their influence on measures of BHI. In the future, respiratory activity should also be added to the physiological measurements of BHI studies (e.g., with a respiration belt).

Finally, the inclusion of (e.g., behavioral or eye movement-related) measures of attention could support the interpretation of emotion-related modulations of attention (as reflected in changes in vagal cardioregulation and parieto-occipital alpha power).

### 5. Conclusion

We replicated previous findings from classical studies in a more naturalistic virtual reality setting, confirming the link between emotional arousal, heart activity, brain oscillations and - albeit less consistently - brain-heart interactions. Our analysis demonstrates that combining measures of heart and brain activity provides insights beyond what can be learned from studying each modality in isolation. Finally, this work illustrates how VR paired with multimodal physiological recordings can be a valuable approach for simultaneously studying multiple components of emotions. Taken together, our results suggest that to better understand affective processes, we must consider the heart alongside the brain, as both play integral roles in emotion.

## Impact statement

Our study shows the value of pairing immersive virtual reality with multimodal physiological recordings for simultaneously studying multiple components of emotions in naturalistic settings. The findings increase our understanding of the link between subjective experience and physiological rhythms not only in the brain but also in the heart.

## Author contributions

A.F.: Conceptualization; Formal analysis; Investigation; Methodology; Project administration; Software; Validation; Visualization; Writing – original draft; Writing – review & editing F.K.: Conceptualization; Data curation; Resources; Writing – review & editing S.M.H.: Conceptualization; Data curation; Resources; Writing – review & editing A.M.: Conceptualization; Data curation; Resources; Writing – review & editing V.V.N.: Conceptualization; Methodology; Writing – review & editing A.V.: Conceptualization; Methodology; Supervision; Writing – review & editing M.G.: Conceptualization; Funding acquisition; Methodology; Project administration; Supervision; Writing – review & editing

## Data and code availability

We did not obtain participants’ consent to release their individual data. Since our analyses focus on the single-subject level, we have only limited data which are sufficiently anonymized (e.g., summarized or averaged) to be publicly shared. All code used for all analyses and plots are publicly available on GitHub at https://github.com/afourcade/evrbhi.

## Acknowledgements

This research was supported by the Max Planck Dahlem Campus of Cognition (MPDCC) and funded by the Max Planck Society - Fraunhofer-Gesellschaft cooperation (project “NEUROHUM”) and the German Federal Ministry of Education and Research (BMBF grant 13GW0488).

## Competing interests

The authors declare no competing interests.

## Notes

### Competing Interest Statement

The authors have declared no competing interest.

### Summary of Updates

Added Ratings section in Results Supplements updated

