## Supplements for "Linking brain-heart interactions to emotional arousal in immersive virtual reality"

### Supplementary Material

**Figure S1.** Experimental setup for the rating phase. A. Participants were rating their emotional arousal during a replay of their experience, with a vertical rating bar visible on the right side of their visual field of view. B. Dial used to rate emotional arousal, ranging from low (0) to high (50) in steps of 1.

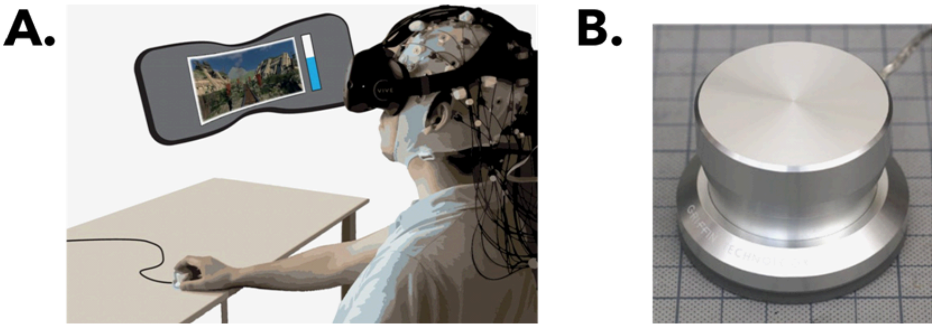

**Figure S2.** Emotional arousal ratings for each participant (colored lines). Black line: mean across participants; vertical lines (light grey): beginning and end of the break. A. With head movement (mov) condition. B. Without head movement (nomov) condition.

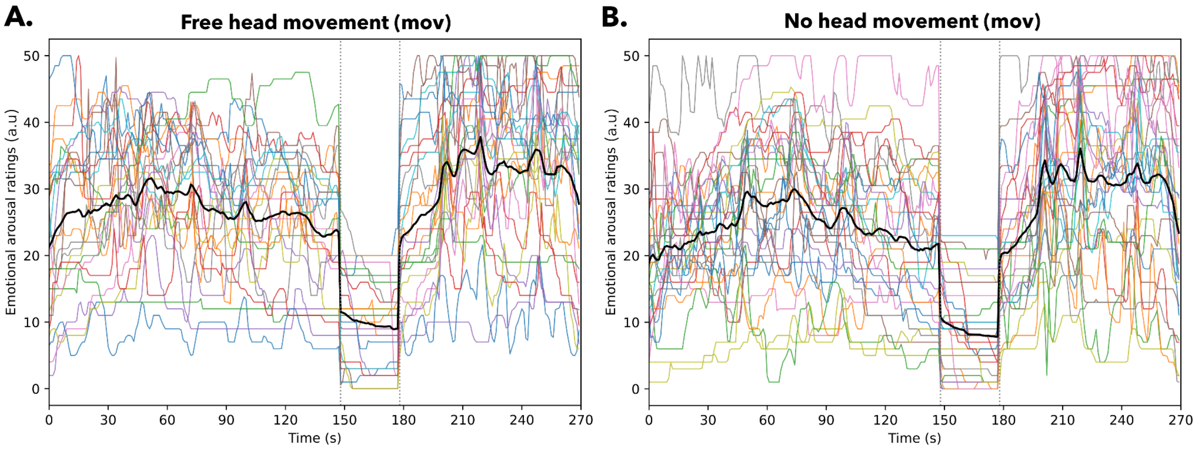

**Figure S3.** Interbeat intervals (IBI) and Low-Frequency Heart Rate Variability (LF-HRV). **A.** No significant effects of arousal ($F(1, 28.3) = 0.02$, $p = .896$), head movement ($F(1, 25.5) = 0.05$, $p = .817$) nor interaction ($F(1, 26.3) = 0.02$, $p = 0.877$) on heart rate. **B.** Significant effect of arousal $F(1, 26.9) = 9.5$, $p = .005$) on LF-HRV. No significant effects of movement ($F(1, 25.5) = 2.2$, $p = .152$) nor interaction ($F(1, 26.0) = 0.5$, $p = .492$). Post-hoc pairwise comparisons of the estimated marginal means revealed significantly lower LF-HRV during higher arousal in the free head movement condition (mov; $t(25.1) = -3.2$, $p = .004$), but not in the without head movement condition (nomov; $t(28.0) = -1.6$, $p = .124$).

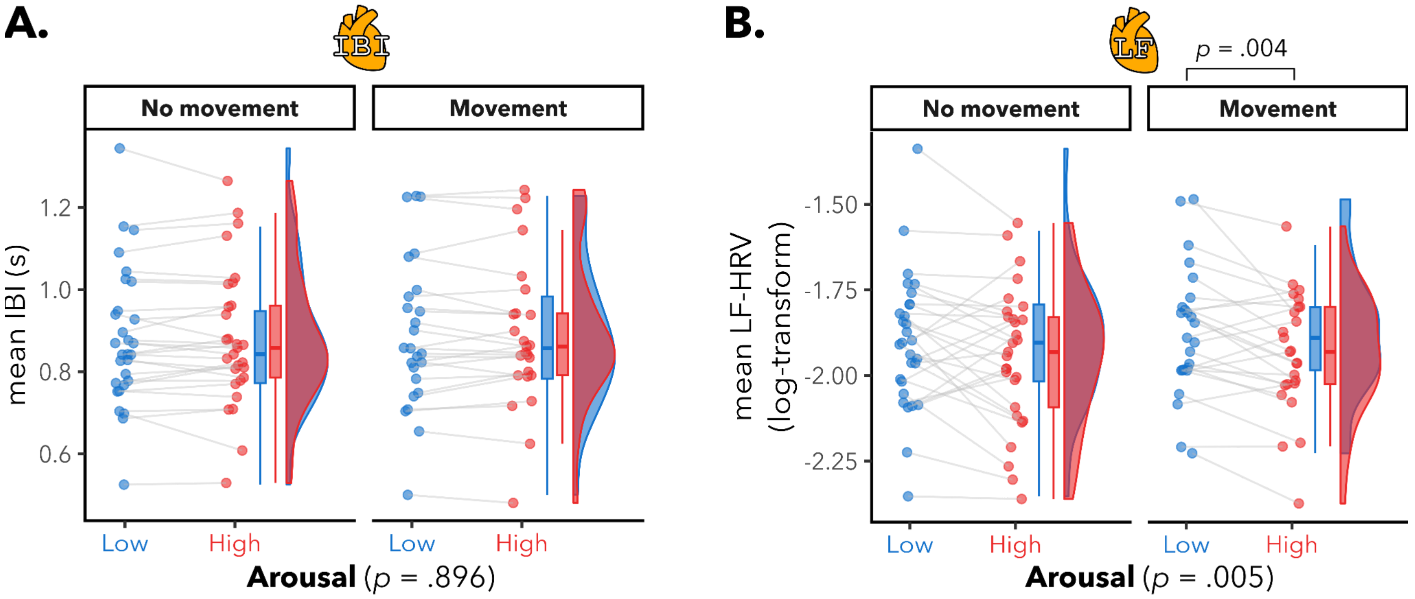

**Figure S4.** Mean time-series across participants of interbeat intervals (IBI), low-frequency heart rate variability (LF-HRV), high-frequency heart rate variability (HF-HRV), α power (averaged across ROI), brain-to-heart coupling coefficient (α → HF-HRV; averaged across ROI), heart-to-brain coupling coefficient (HF-HRV → α; averaged across ROI) and emotional arousal ratings, for the condition without head movement (nomov).

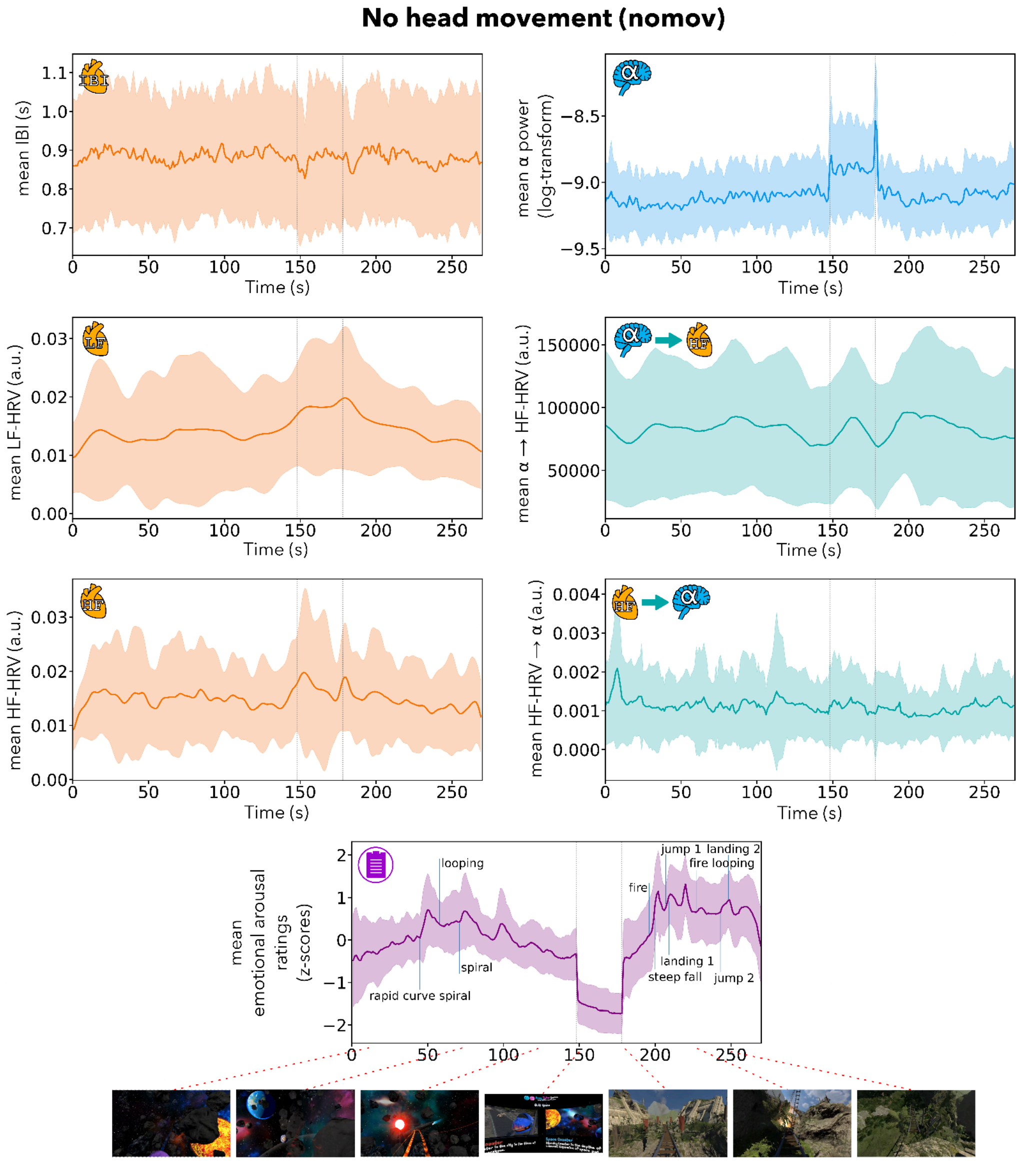

**Figure S5.** Other frequency bands power - Brain/EEG

Topographic plots of t-values showing significant electrodes ($p_{uncorrected} < .05$; $p_{FDR} < .05$ in Bold) from paired t-test of the mean HA vs. mean LA for each participant (averaged over the head movement conditions), at each electrode, for each EEG frequency band.

**A.** No electrodes with significant HA vs. LA differences in delta power. **B.** Lower theta power during higher arousal in 17 electrodes (Fp1, Fp2, Fz, F3, FC1, FC2, FC6, C3, C4, CP1, CP5, P4, P7, TP9, TP10, O1 and O2). **C.** Lower alpha power during higher arousal in 23 electrodes (Fp1, Fp2, Fz, F3, F4, FC1, FC2, FC6, Cz, C4, CP1, CP5, CP6, Pz, P3, P4, P7, P8, TP9, TP10, T8, O1 and O2). **D.** Lower beta power during higher arousal in 6 electrodes (FC2, C4, CP1, Pz, P3 and P4). **E.** Higher gamma power during higher arousal in electrode O2.

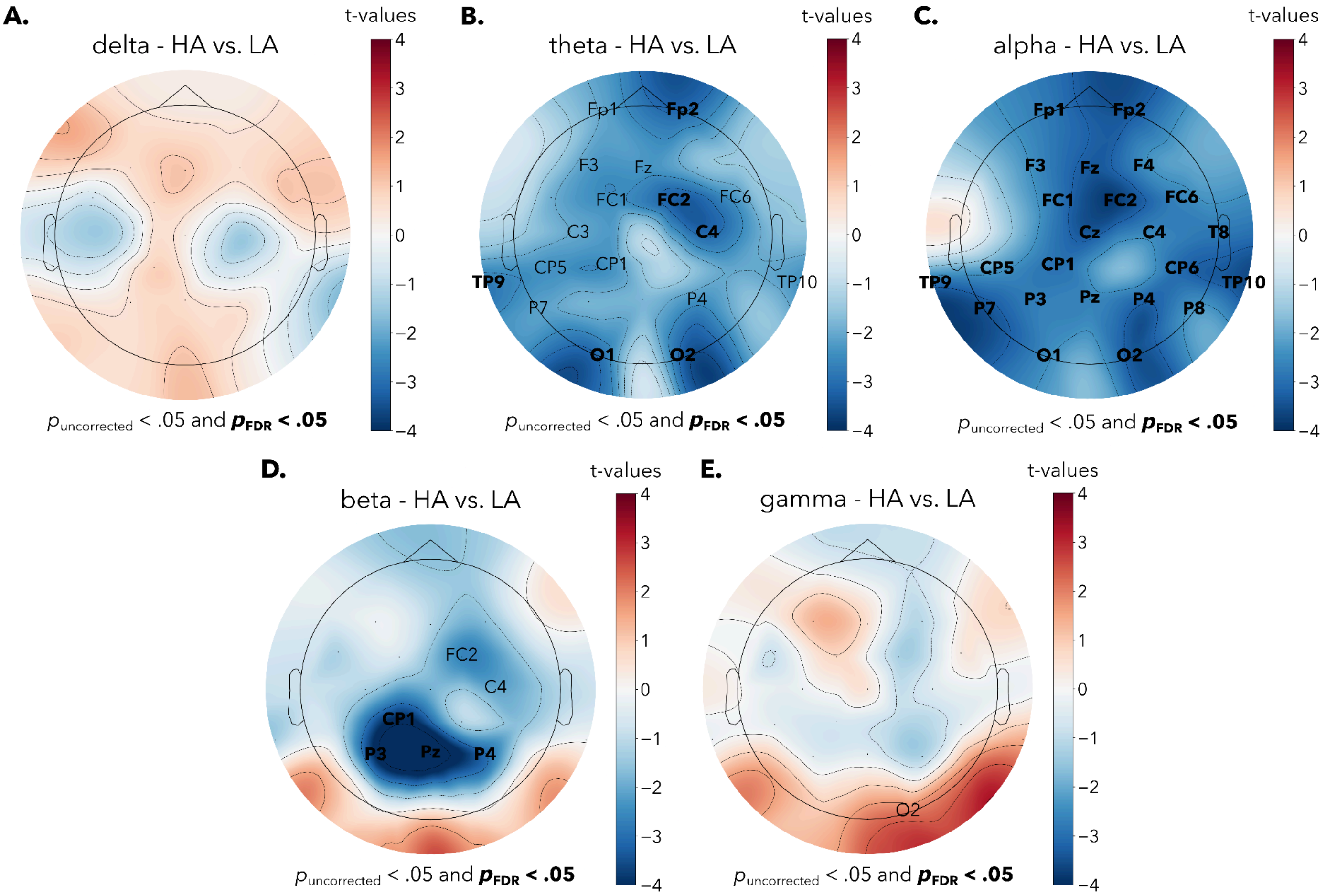

**Figure S6.** Other frequency bands power - Brain-Heart

Topographic plots of t-values showing significant electrodes ($p_{uncorrected} < .05$; $p_{FDR} < .05$ in Bold) from paired t-test of the mean HA vs. mean LA for each participant (averaged over the head movement conditions), at each electrode, for each directional Brain-Heart Interaction (BHI) coupling coefficient.

For HF-HRV → δ, the EEG electrode O1 showed lower BHI coupling coefficients during higher arousal. For LF-HRV → δ, the EEG electrode P8 showed higher BHI coupling coefficients during higher arousal. For HF-HRV → α, the EEG electrode P3 showed lower BHI coupling coefficients during higher arousal. For HF-HRV → β, the EEG electrodes Pz and CP2 showed lower BHI coupling coefficients during higher arousal. For HF-HRV → γ, the EEG electrode O2 showed higher BHI coupling coefficients during higher arousal. For LF-HRV → γ, there were 12 EEG electrodes (Fp2, Fz, FC5, CP5, P3, P8, T7, TP9, TP10, Oz, O1, O2) with higher BHI coupling coefficients during higher arousal. For γ → HF-HRV, there were 17 EEG electrodes (Fp1, F3, FC5, FC6, C3, CP5, CP6, P3, P4, P7, T7, T8, TP9, TP10, Oz, O1, O2) with lower BHI coupling coefficients during higher arousal. Furthermore, concerning the gamma band, when looking at the interception (FC5, CP5, P3, T7, TP9, TP10, O1, Oz, O2) of both significant electrodes for the ascending (LF-HRV → γ) and descending (γ → HF-HRV) directions, both effects were still present (LF-HRV → γ: $t(28) = 2.8$; $p = .008$; γ → HF-HRV: $t(28) = -2.7$; $p = .012$).

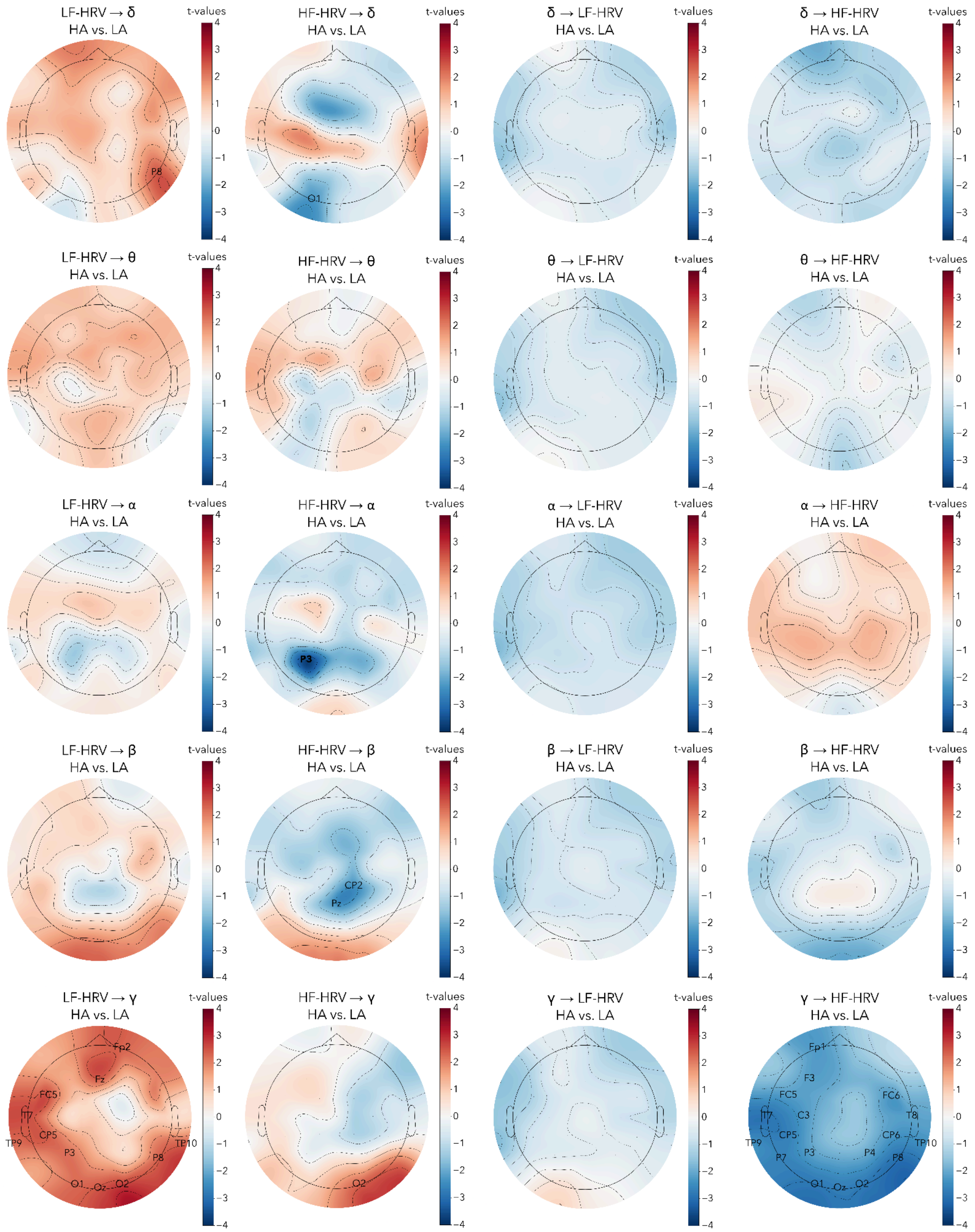


$p_{uncorrected} < .05$ and $\boldsymbol{p_{FDR} < .05}$
